# Activator–promoter compatibility in mammals: a CpG-Island-specific co-activator directly bridges transcription factors to TFIID

**DOI:** 10.64898/2025.12.28.696747

**Authors:** Filip Nemčko, Kevin Sabath, Michaela Pagani, Sebastian Kleinfercher, Clemens Plaschka, Alexander Stark

## Abstract

Transcription from CpG island (CGI) promoters controls the expression of two-thirds of mammalian genes, yet despite their prevalence, it remains unknown whether CGI-specific co-activators with intrinsic specificity (i.e. compatibility) for these promoters exist or by what mechanisms they might function. Here, we perform proteome-wide functional screens to identify more than fifty transcriptional activators that are intrinsically specific to CGI promoters, establishing that promoter-class-specific activators are a widespread feature of mammalian gene regulation. Among these, we identify Host Cell Factor 1 (Hcfc1) as the founding member of CGI-specific co-activators. Hcfc1 is essential for the expression of thousands of CGI-promoter-driven genes and acquires CGI-specificity through a two-step mechanism: CGI-associated transcription factors recruit Hcfc1 through its Kelch domain, and Hcfc1 in turn directly engages the general transcription factor TFIID through a dedicated activation domain. The Hcfc1-TFIID interaction overcomes a key rate-limiting step for CGI promoter initiation, TFIID recruitment, thereby directly enabling transcription. Hcfc1 thus functions as a promoter-class-specific bridge between CGI-bound transcription factors and the general transcription machinery, analogous to Mediator but with intrinsic promoter specificity. Together, we uncover a dedicated activation pathway for CGI promoters, reveal a fundamental mechanistic difference in transcription activation between promoter classes in mammals, and establish co-activator-promoter compatibility as a core principle in mammalian gene regulation.

## Introduction

The transcription of protein-coding genes is a fundamental step in eukaryotic gene expression. Transcription initiates at the 5’ end of genes within specific DNA sequences termed promoters^1^. Studies on transcription initiation have largely focused on the class of TATA-box promoters, where transcription begins from a single dominant site around 30 nucleotides downstream of the TATA-box motif^2^. In mammals, however, approximately two-thirds of all promoters constitute a distinct promoter type rich in CpG dinucleotides (**Fig. 1a**). These CpG island (CGI) promoters include the majority of housekeeping and key developmental genes and differ from TATA-box promoters in important aspects, including dispersed transcription start sites and a tightly positioned +1 nucleosome with strongly phased downstream nucleosomes^1,3,4^. This suggests distinct mechanistic requirements for transcription initiation between these two promoter classes.

**Fig. 1:**
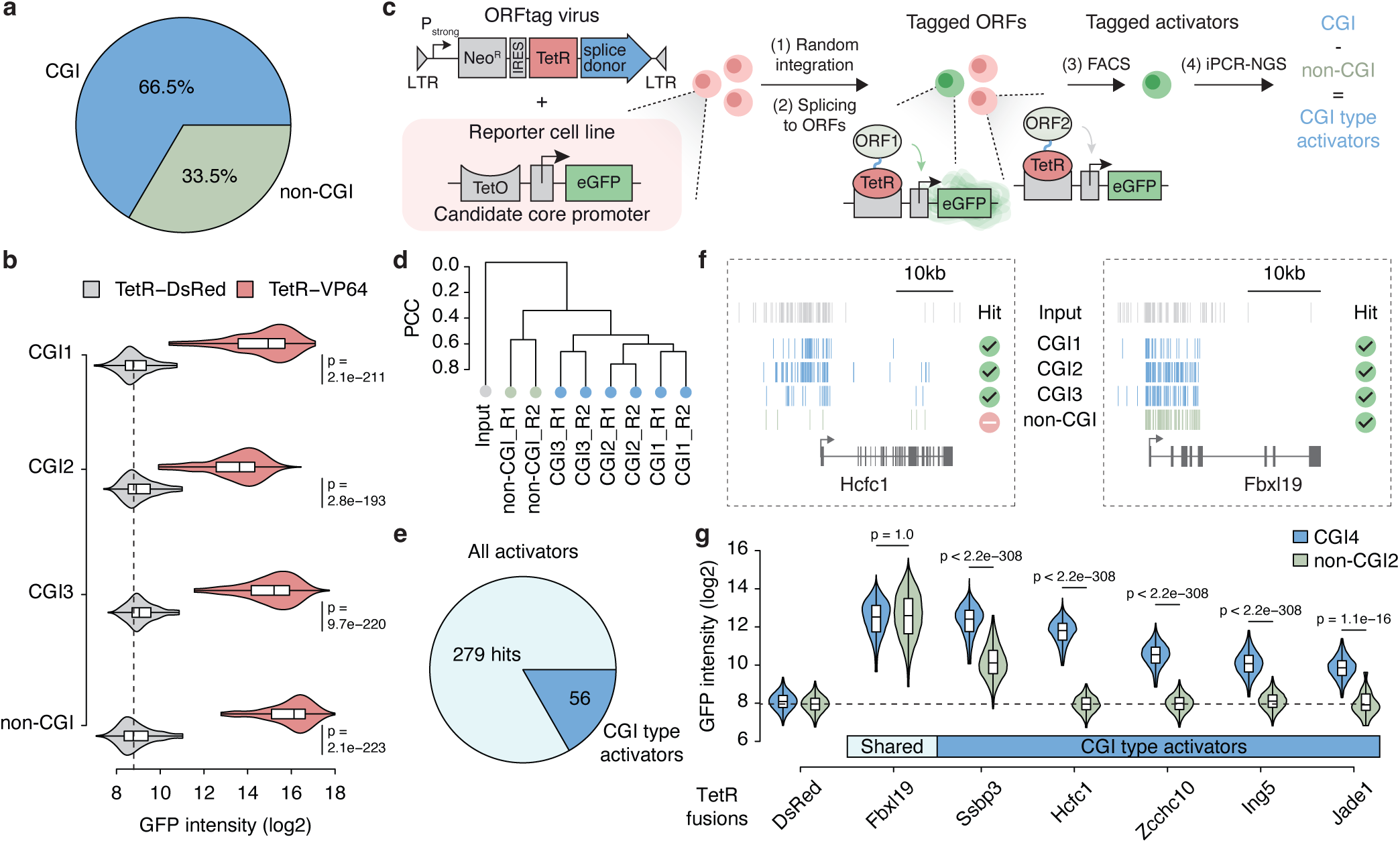
ORFtag screens identify Hcfc1 as a potent CGI-specific transcriptional activator. **a**, Proportion of mouse promoters that overlap with CpG islands (CGIs). **b**, GFP intensity in reporter cell lines stably expressing TetR-DsRed (control) or TetR-VP64, measured by flow cytometry. The sample size was 700 cells. Statistical significance was assessed using a one-sided Wilcoxon test. Box plots indicate the median (central line), 25^th^-75^th^ percentiles (box), and 1.5x interquartile range (whiskers). **c**, Schematic of the ORFtag approach. Pstrong, strong promoter; eGFP, enhanced GFP; iPCR-NGS, inverse PCR followed by next-generation sequencing; FACS, fluorescence-activated cell sorting. **d**, Dendrogram showing Pearson Correlation Coefficients (PCC) between merged Input (grey) and sorted sample replicates (blue, green). Input replicates exhibited high PCC (≥ 0.91). **e**, Number of ORFtag hits that activated all three CGI promoters but not the non-CGI control. **f**, Genome browser views of ORFtag integration sites (vertical lines, correct orientation only) at the Hcfc1 (left) and Fbxl19 (right) loci, before (Input; grey) and after FACS selection. **g**, GFP intensity was measured by flow cytometry in reporter cell lines stably expressing the indicated full-length proteins fused to TetR. The sample size was 20,000 cells. Statistical analysis and box plots are as described in (**b**).

The differences between promoter classes might be explained through their regulation by distinct transcription co-activators. Some co-activators *might merely be recruited in a promoter-type-specific manner while others might* possess protein-intrinsic properties that allow them to activate specific promoter types but not others, *even if recruited to both promoter types*. Such biochemical *co-activator–promoter compatibilities* can be revealed by forced recruitment of co-activators to promoters via heterologous DNA-binding domains^5^. Examples of compatibility have been reported in *Drosophila*^6,7^, but whether such compatibilities exist in mammals, especially with a preference for CGI promoters, remains less clear. The nature of regulatory mechanisms and co-activators for CGI promoter activation, the most prevalent promoter class in mammals, thus remains a major knowledge gap in our understanding of transcription regulation in mammals, including humans.

Here, we identify 56 CGI-specific activators, including Host Cell Factor 1 (Hcfc1) and demonstrate that Hcfc1 is both sufficient to activate CGI promoters upon forced recruitment and required for thousands of active CGI promoters. Hcfc1 has been previously reported as a conserved transcriptional co-regulator that is recruited to chromatin through interactions with transcription factors (TFs) containing Hcfc1-binding motifs (HBMs)^8,9^. These TFs, including Thap11, Myc and others^10–12^ are enriched at active CGI promoters consistent with observations that Hcfc1 is frequently detected at such promoters in human cells^13^. However, how Hcfc1 functionally contributes to CGI promoter activation, and by what mechanism, has remained unknown.

We further identify TFIID recruitment as a key rate-limiting step in CGI promoter activation and show that Hcfc1 directly bridges CGI-specific TFs to TFIID, thus directly promoting transcription. By revealing a dedicated co-activator that physically links CGI-bound TFs to TFIID, our work uncovers a previously unrecognized pathway for initiating transcription at CGI promoters, reveals a fundamental mechanistic difference between promoter classes in mammals, and establishes co-activator-promoter compatibility as a core principle in mammalian gene regulation.

## Results

### ORFtag identifies Hcfc1 as a CGI-specific co-activator

To systematically identify transcriptional activators with intrinsic compatibilities towards CGI promoters, we performed functional screens in mouse embryonic stem cells (mESCs) based on force-recruiting candidate proteins to stably chromosomally integrated eGFP reporter genes, bearing three different minimal CGI promoters from the NCAPG, ELAC2, and DYNC1LI1 genes (**Extended Data Fig. 1a**). For each promoter, we added an upstream tetO array and a downstream eGFP open-reading frame and individually integrated the resulting eGFP reporter constructs into a constant landing site on chromosome 15 as described before^14^. All three eGFP reporters were inactive in the absence of an activator but were strongly induced by force-recruiting the strong activator TetR-VP64^15^. The low basal and high induced activities were similar to a previously published eGFP reporter bearing the non-CGI minimal promoter of the MYLPF gene^14^ (**Extended Data Fig. 1b**).

We next performed proteome-wide screens for transcriptional activators in each of the three CGI-reporter cell lines using ORFtag^14^, two independent replicates each. Briefly, we infected 150 million cells per replicate with a retrovirus bearing a TetR-ORFtag cassette to tag endogenous protein-coding exons with the Tetracycline Repressor (TetR) DNA-binding domain to recruit the tagged proteins to the tetO array of the respective eGFP reporter (**Fig. 1c**). We isolated GFP-positive cells by fluorescence-activated cell sorting (FACS) and identified tagged candidate proteins via an enrichment of viral insertion sites in GFP-positive cells compared to the non-sorted input sample^14^.

The three CGI screens showed strong consistency between replicates and clustered separately from a previously published ORFtag screen with the MYLPF non-CGI promoter (**Fig. 1d, Extended Data Fig. 1b**). To identify CGI-specific activators, we required that candidate proteins activated all three CGI promoters (false discovery rate (FDR) < 0.1%, log2 Odds Ratio ≥ 1) but not the non-CGI control (FDR < 5%, log2 Odds Ratio ≥ 1), which defined 56 high-confidence CGI-specific activators (**Fig. 1e, Supplementary Table 1**). Raw ORFtag data for Hcfc1 illustrate this specificity: in contrast to integrations tagging a general activator like Fbxl19, which were enriched for all four screens, Hcfc1-tagging integrations were enriched only in the three CGI screens (**Fig. 1f**).

To validate the screening hits, we individually cloned TetR-fusion constructs for five CGI-specific candidates and one shared activator and force recruited them to independent genomically integrated CGI and non-CGI reporters, derived from CHMP3 and SPINK5 genes respectively, promoters that have not been used for the screens (**Fig. 1a and c**). All five CGI-specific candidates preferentially activated the CGI reporter and with four candidates (Hcfc1, Zcchc10, Ing5, and Jade1) exclusively activating the CGI reporter, i.e. being CGI specific (**Fig. 1g**). Among these, Hcfc1 displayed the most potent and specific activity, altogether activating four out of four CGI but not the two non-CGI promoters.

Originally discovered as a host factor for herpes simplex virus (HSV) and the cellular target of HSV VP16^12^, Hcfc1 is known to be an essential co-factor for cell proliferation and the transcription of Myc-driven housekeeping genes^16^. Moreover, its localization to CGI promoters in HeLa cells^13^ makes it a promising candidate for a first CGI-specific co-factor with exclusive CGI compatibility. We therefore selected Hcfc1 to investigate the mechanistic basis of a potential CGI-specific cellular role and an exclusively CGI-promoter compatible function.

### Hcfc1 localizes to CGI promoters in mESCs

A CGI-specific function of Hcfc1 would predict its localization to CGI promoters also in mESCs. To test this, we mapped its genome-wide occupancy using Cleavage Under Targets and Release Using Nuclease (CUT&RUN) in mESCs (**Fig. 2a**). Consistent with its localization in HeLa cells, 58% (10,905 of 18,651) of Hcfc1 binding sites in mESCs overlapped promoters (**Fig. 2a and b**), of which more than 94% were CGI promoters (**Fig. 2c**). Thus, Hcfc1 indeed exhibits a strong preference for binding to CGI promoters^13^.

**Fig. 2:**
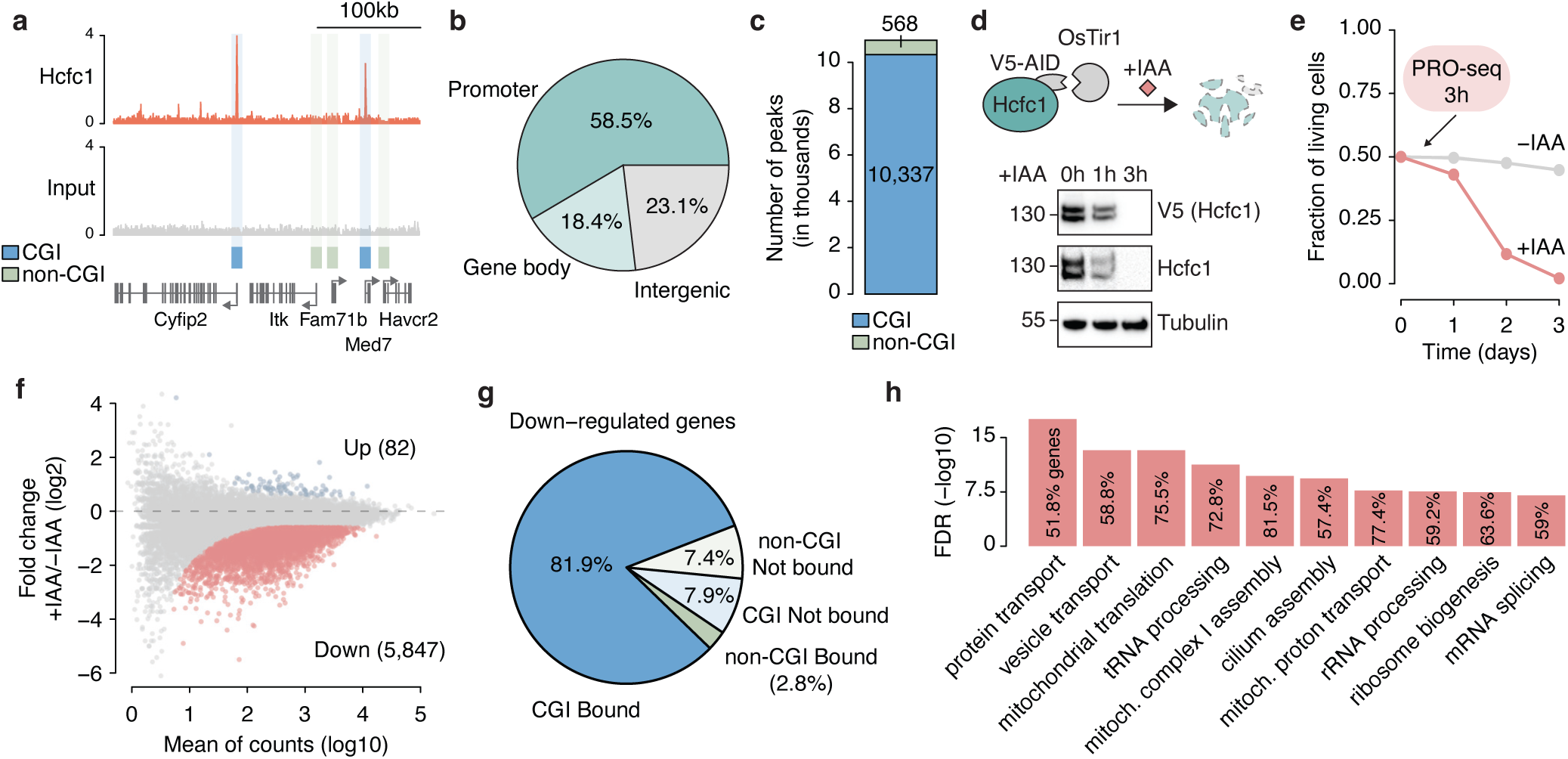
Hcfc1 binds thousands of CGI promoters and is required for their transcription. **a**, Genome browser view of Hcfc1 CUT&RUN signal. CGI (blue) and non-CGI (green) promoters are marked for reference. **b**, Genomic distribution of Hcfc1 peaks (n=18,651), showing the proportion located at promoters (±1 kb from the gene start), within gene bodies, and in intergenic regions. **c**, Breakdown of promoter-associated peaks from (**b**), showing the number overlapping with CGI versus non-CGI promoters. **d**, Schematic of the rapid Hcfc1 depletion using the AID system. The western blot is representative of three independent experiments and shows Hcfc1 loss after 3 h of indole-3-acetic acid (IAA) treatment. **e**, Time-course analysis of cell viability, comparing cells with Hcfc1 present (-IAA, grey) to those where Hcfc1 is depleted (+IAA, red). Data are from three biological replicates. **f**, MA plot of PRO-seq log2 fold changes (log2FC) in transcription after 3 h of Hcfc1 depletion. Significantly up- (blue) and down-regulated (red) genes are highlighted (FDR < 0.05, log2FC > log2(1.5)). Data are from two biological replicates. **g**, Classification of down-regulated genes based on Hcfc1 promoter binding status (bound or unbound) and promoter type (CGI or non-CGI). **h**, Top 10 enriched GO terms for biological processes among Hcfc1-dependent genes, based on a Fisher’s exact test (FDR-adjusted P-values).

### Hcfc1 is essential for transcription from thousands of CGI promoters

While Hcfc1 was sufficient to activate CGI core promoters upon recruitment and localized to CGI promoters in the genome, we next asked whether Hcfc1 is necessary for the activity of endogenous CGI promoters. To assess the direct transcriptional impact of Hcfc1 loss while minimizing secondary effects, we employed rapid protein depletion using an auxin-inducible degron (AID) system in mES cells expressing the Tir1 E3 ligase^17^ in which Hcfc1 is homozygously tagged. Auxin treatment rapidly depleted Hcfc1 protein within 3 hours (**Fig. 2d**) and, as expected for an essential gene^18,19^, prolonged Hcfc1 depletion resulted in cell death after two to three days (**Fig. 2e**). To capture the immediate transcriptional effects, we measured nascent transcription changes by precision nuclear run-on sequencing (PRO-seq) 3 hours after Hcfc1 depletion (**Extended Data Fig. 2b, Supplementary Table 2**). This revealed significant down-regulation (> 1.5-fold change, false discovery rate < 0.05) of over 5,800 genes (**Fig. 2f**), of which the vast majority (89.8%) had CGI promoters, and of which most (91.2%) had Hcfc1 bound at their promoters (**Fig. 2g**). Gene Ontology (GO) analysis showed that the Hcfc1-dependent genes were highly enriched for terms related to essential housekeeping functions (e.g. protein transport, mitochondrial translation, tRNA processing) (**Fig. 2h**). This aligns with the known association of CGI promoters with ubiquitously expressed genes^4^. Together, the data so far show that Hcfc1 recruitment is sufficient to activate CGI but not non-CGI promoters and that it binds to and is necessary for the activity of thousands of endogenous CGI promoters, suggesting that it functions as a dedicated transcriptional co-activator for this promoter class and that it might constitute a founding member of CGI-promoter-specific activators. We therefore set out to determine the mechanisms of its recruitment to CGI promoters and its ability to activate transcription at CGI promoters.

### Hcfc1 is targeted to CGI promoters via its Kelch domain

Hcfc1 lacks a dedicated DNA-binding domain, suggesting that its recruitment to CGI promoters is mediated through protein-protein interactions. We used TurboID proximity labelling in mES cells to capture Hcfc1’s proximal interactome (**Extended Data Fig. 3a, Supplementary Table 3**). This recovered known interactions with co-activators (CoAs) such as COMPASS-like (SET1/MLL) complexes^20^, NSL complex^21^ and various TFs, which constituted 44% of all significant hits (56/127, **Fig. 3a**). Moreover, analysis of the detected TFs’ known DNA-binding motifs revealed a significantly higher GC content compared to those of other TFs expressed in mES cells. Similarly, several of the detected co-activators contain CxxC domains that recognize CpG dinucleotides^22^. These results are consistent with a model in which CxxC-containing co-activators and TFs that recognize CpG-rich sequences would recruit Hcfc1 to CGI promoters (**Fig. 3b**).

**Fig. 3:**
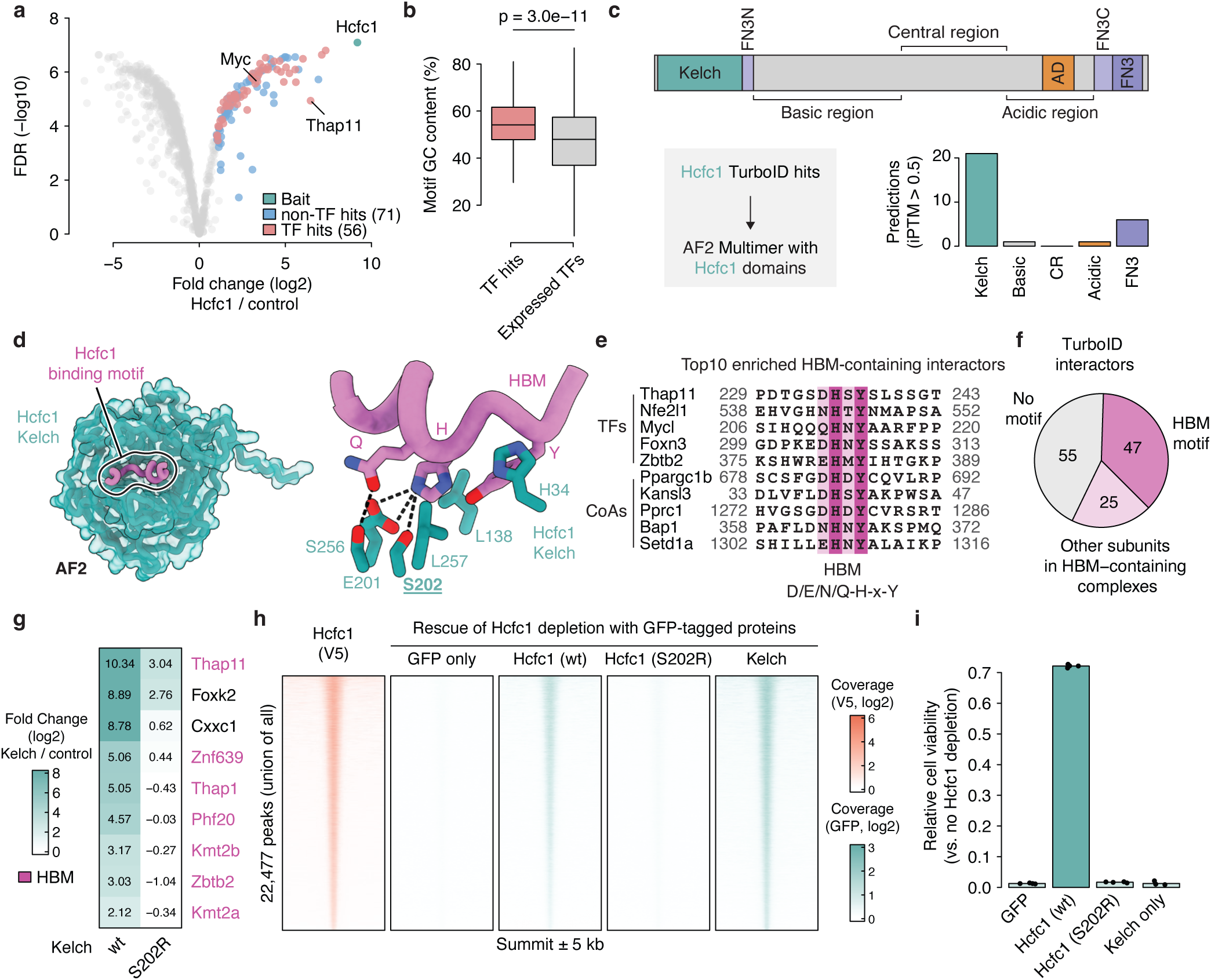
The Hcfc1 Kelch domain is targeted to CGI promoters via HBM-containing transcription factors. **a**, Volcano plot of proteins identified by Hcfc1 TurboID versus a control TurboID. Significantly enriched proteins (log2FC > 1, FDR < 0.05) are coloured. Data are from three biological replicates. **b**, Mean GC content of motifs for Hcfc1-interacting TFs compared to motifs for all expressed TFs. Statistical significance was assessed using a one-sided Wilcoxon test. Box plots indicate the median (central line), 25^th^-75^th^ percentiles (box), and 1.5x interquartile range (whiskers). **c**, Top: Domain organization of Hcfc1. FN3, Fibronectin type III domain; FN3N/C, split Fibronectin type III domain; AD, activation domain. Bottom: Schematic (left) and results (right) of an *in silico* screen using AF2 to identify direct Hcfc1 interactors. Predictions with an average interface predicted TM (iPTM) score > 0.5 are shown. **d**, AF2 model of the Hcfc1 Kelch domain (green) in complex with an HBM-containing interactor (pink), shown in overview (left) and close-up (right). Key interacting residues are labelled, hydrogen bonds are indicated by dashed lines, and the residue mutated in this study is underlined. **e**, Sequence alignment of HBMs from Hcfc1 interactors identified by TurboID. Residues with complete conservation (pink) and > 70% similarity (light pink) are highlighted. The consensus HBM sequence is shown below. **f**, Proportion of Hcfc1 TurboID interactors that contain an HBM or belong to complexes with an HBM-containing protein. **g**, Heatmap of protein enrichment from IP-MS of the wild-type or S202R mutant Hcfc1 Kelch domain versus a control. Data are from three biological replicates. The top 9 TFs significantly enriched in the wild-type sample are shown, HBM-containing proteins are highlighted in pink. **h**, Heatmaps of CUT&RUN signal for endogenously tagged Hcfc1 (orange) and exogenously expressed GFP-tagged rescue constructs (green). Heatmaps display signal across the union of all peaks, with each row centred on the peak summit over a ±5kb window. **i**, Relative viability of cells expressing the rescue constructs from (H). Results show the fraction of viable cells after 3 days of endogenous Hcfc1 depletion (+IAA), normalized to untreated controls (-IAA) for each line (n=4).

We next sought to define the molecular basis of Hcfc1-TF interactions. We used AlphaFold2-Multimer (AF2) to predict the interfaces between Hcfc1 and its putative interactors^23,24^. This identified the N-terminal Kelch domain as the primary interaction hub (**Fig. 3c**), consistent with earlier reports^8,9^. Manual inspection of the predicted models revealed a minimal motif, D/E/N/Q-H-x-Y, that docks into a conserved pocket on the Kelch domain surface (**Fig. 3d, Extended Data Fig. 3b-d**). This motif was prevalent in the TurboID hits and corresponds to the previously identified Hcfc1-binding motif (HBM^8,12^). In total, we identified 72 TurboID hits that either directly contain an HBM or are in complexes with an HBM-containing protein. Most interactors are linked to transcription, as 83% (60/72) of these factors are TFs or CoAs (**Fig. 3e and f**). To functionally test if the Kelch-HBM interface mediates Hcfc1 binding, we designed a point mutant in the Kelch domain (S202R) to disrupt the predicted interaction (**Fig. 3d**). Immunoprecipitation followed by mass spectrometry (IP-MS) confirmed the prediction: while the wild-type Kelch domain robustly enriched HBM-containing interactors, including co-activators and TFs, the single S202R mutation strongly reduced or completely abolished these interactions (**Fig. 3g, Extended Data Fig. 3e and f, Supplementary Table 3**).

To determine the function of this interface in recruiting Hcfc1 to the genome, we used the AID system to deplete endogenous Hcfc1 in mES cells while expressing rescue constructs. CUT&RUN analysis demonstrated that Hcfc1’s Kelch domain alone was sufficient to occupy its native binding sites. However, Kelch alone failed to rescue cell viability, leading to cell death after two to three days. Furthermore, full-length Hcfc1 bearing the S202R mutation failed to be recruited to its own endogenous sites and caused cell death within the same timeframe. In contrast, wild-type Hcfc1 successfully restored recruitment to its endogenous binding sites and rescued the lethality of Hcfc1 depletion. These findings establish that the Kelch-HBM interface is both necessary and sufficient for Hcfc1 recruitment to CGI promoters (**Fig. 3h and i, Extended Data Fig. 4**).

### Hcfc1 recruits TFIID via a CGI-specific activation domain

To define the mechanism of Hcfc1-mediated CGI-specific transcription activation, we first sought to identify the domain responsible for its activation function. Guided by previous reports, we identified an activation domain (AD) within the C-terminal acidic region (residues 1600-1730)^25^. The isolated Hcfc1-AD recapitulated the strict CGI specificity of full-length Hcfc1, activating the CGI-but not the non-CGI reporter (**Fig. 4a**), thus constituting a first CGI-specific AD.

**Fig. 4:**
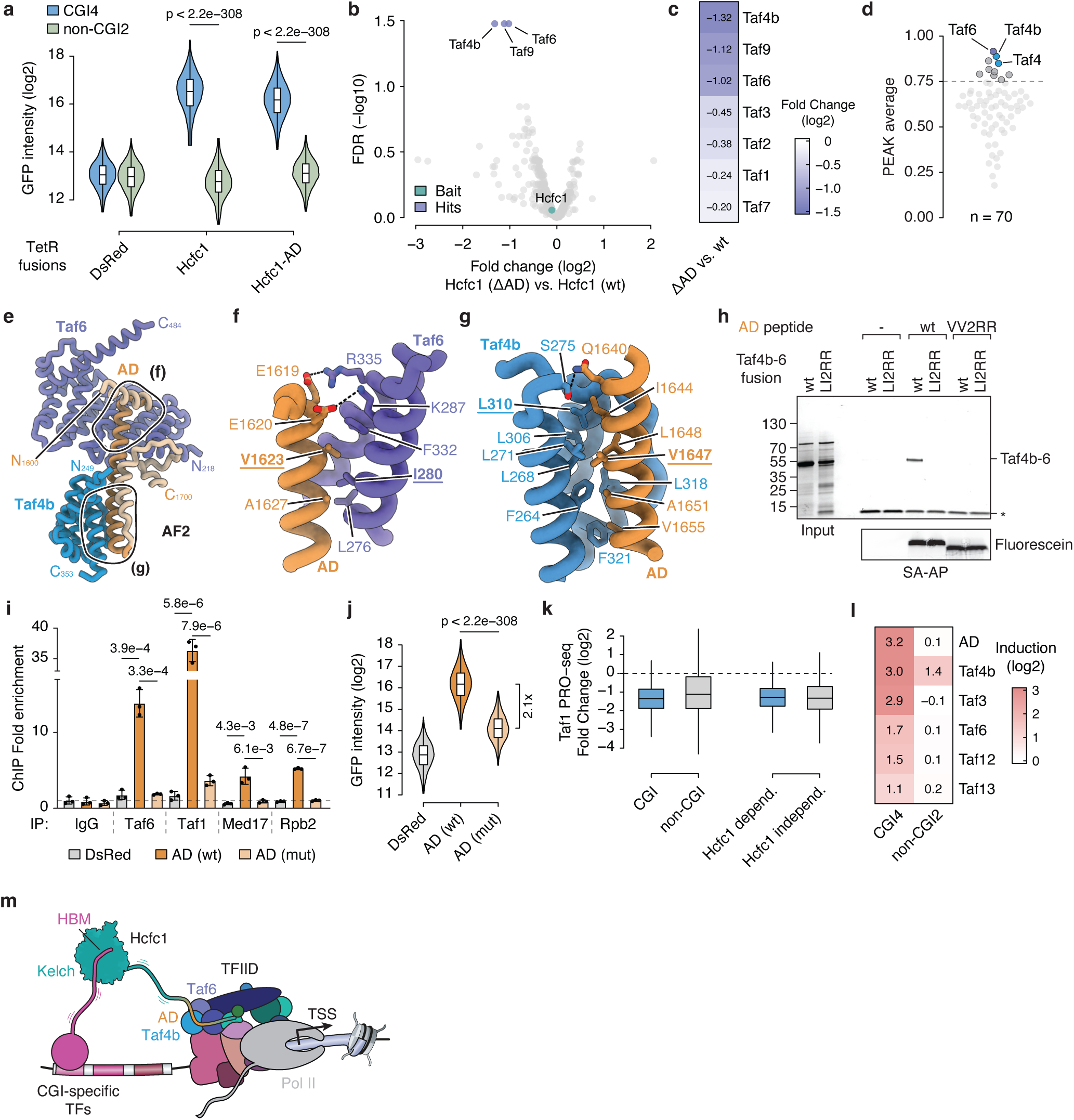
A CGI-specific activation domain of Hcfc1 recruits the TFIID complex ab. Mapping the AD of Hcfc1. GFP intensity was measured by flow cytometry in reporter cell lines stably expressing the indicated constructs fused to TetR. Statistical significance was assessed using a one-sided Wilcoxon test. Box plots indicate the median (central line), 25^th^-75^th^ percentiles (box), and 1.5x interquartile range (whiskers). **b**, Volcano plot comparing the TurboID profiles of Hcfc1 ΔAD versus wild-type Hcfc1. Data are from three biological replicates. Significantly depleted proteins in the Hcfc1 ΔAD sample (log2FC > 1, FDR < 0.05) are coloured. **c**, Heatmap showing the relative depletion of top 7 TFIID subunits from the TurboID comparison in (**b**). **d**, AF2 *in silico* screen to identify direct interactors of the Hcfc1-AD. Predictions with an average PEAK score > 0.75 are highlighted. **e**, AF2 model of the Hcfc1-AD (orange) interacting with Taf4b (blue) and Taf6 (violet). **f**,**g**, Close-up views of the interaction interfaces between the Hcfc1-AD and Taf6 (**f**) and Taf4b (**g**). Interacting residues are labelled, hydrogen bonds are shown as dashed lines, and residues mutated in this study are underlined. **h**, Co-affinity purification (co-AP) assay testing the interaction between wild-type (wt) or mutant (VV2RR) Hcfc1-AD core peptides and a wt or mutant (LI2RR) Taf4b-Taf6 fusion protein. Results are shown by Coomassie-stained SDS-PAGE. Equal bait loading was confirmed by fluorescein intensity. Asterisk (*), streptavidin. **i**, ChIP-qPCR analysis of TFIID, Mediator, and Pol II enrichment at a CGI4 reporter promoter. Enrichments are normalized to a negative control locus. Bar graphs display the mean of three biological replicates, error bars representing the standard deviation. Statistical significance was determined using two-sided Student’s t-test. **j**, GFP intensity measured by flow cytometry in reporter cell lines stably expressing the indicated constructs fused to TetR. Statistical analysis and box plots are as described in (**a**). **k**, Boxplot of PRO-seq log2FC in transcription after 3 h of Taf1 depletion for CGI and non-CGI genes, and for Hcfc1-dependent and -independent genes. Box plots are as described in (**a**). Data are from two biological replicates. **l**, Heatmap showing GFP induction (relative to TetR-DsRed) in reporter cell lines stably expressing the indicated TFIID subunits fused to TetR. GFP intensity was measured by flow cytometry. **m**, Summary model for Hcfc1-mediated activation of CGI-promoters. Hcfc1 is recruited to CGI promoters via HBM-containing TFs, and the Hcfc1-AD directly recruits TFIID to drive transcription.

The strict CGI specificity of the Hcfc1-AD led us to hypothesize that it interacts with protein(s) that confer CGI promoter selectivity and/or regulate a rate-limiting step unique to CGI promoters. To identify these interactors, we performed comparative proximity labelling with TurboID, using full-length Hcfc1 versus an Hcfc1-AD deletion mutant (ΔAD) as bait (**Extended Data Fig. 5a, Supplementary Table 3)**. While most interactors, including the Kelch-domain-binding TFs, were enriched to similar levels for both baits, three subunits of the general transcription factor complex TFIID (Taf4b, Taf6, and Taf9) were significantly depleted upon Hcfc1-AD deletion (**Fig. 4b, Extended Data Fig. 5b**).

Other TFIID subunits also showed reduced enrichment compared to full-length Hcfc1 (**Fig. 4c**), suggesting that the Hcfc1-AD might recruit the multi-subunit complex TFIID, which is essential for promoter recognition, recruitment of the transcription machinery, and transcription initiation at RNA polymerase II (Pol II) promoters^26^.

To model the molecular basis of this interaction, we used AF2 to generate predictions between the Hcfc1-AD and components of the transcription pre-initiation complex (PIC), including TFIID (**Fig. 4d**). This analysis revealed a high-confidence trimeric complex, comprising the Hcfc1-AD, Taf4b, and Taf6 – the same TFIID subunits identified by the TurboID experiment (**Fig. 4e-g, Extended Data Fig. 5c and d**). Taf4b is a paralog of the core TFIID subunit Taf4, linked to germ cell transcription programs^27–29^, and Taf6 is a central structural subunit forming multiple interactions within TFIID^26^. Notably, based on known cryo-EM structures, the predicted interaction surfaces on Taf4b and Taf6 are accessible in the context of the entire TFIID complex and the TFIID-bound PIC (**Extended Data Fig. 5E**)^30^.

To test the predicted Hcfc1-Taf4b-Taf6 interactions, we designed a double point mutant in the Hcfc1-AD (V1623R, V1647R), predicted to abolish these interactions (**Fig. 4f and g**). Co-affinity purification using synthetic Hcfc1-AD peptides and a purified Taf4b-Taf6 fusion protein revealed an interaction between Taf4b-Taf6 and the wild-type Hcfc1-AD but not the VV2RR mutant peptide. To further validate this interface, we performed the reciprocal experiment: mutations in Taf4b (L310R) and Taf6 (I280R) also abolished the interaction with the wild-type Hcfc1-AD peptide (**Fig. 4h**). Together, these data demonstrate a direct and specific physical interaction between Hcfc1-AD and the TFIID subunits Taf4b and Taf6.

To determine if the Hcfc1-AD – TFIID interaction leads to the recruitment of TFIID to CGI promoters, we tethered a TetR – Hcfc1-AD fusion protein to the tetO-CGI4-eGFP reporter and performed chromatin immunoprecipitation followed by quantitative PCR (ChIP-qPCR). TetR fused to the wild-type Hcfc1-AD efficiently recruited the TFIID subunits Taf6 and Taf1, as well as Mediator subunit Med17 and the Pol II subunit Rpb2, indicating PIC assembly at the promoter. In contrast, TetR fused to the VV2RR mutant Hcfc1-AD lead to little or no recruitment of any of the four tested factors (**Fig. 4i, Extended Data Fig. 6a**), and – as expected – did not activate reporter transcription (**Fig. 4j**). Together, these results validate that the Hcfc1-AD – Taf4b/6 interface is the critical link for recruiting the transcription machinery and activating CGI promoters.

These results suggest that Hcfc1-AD achieves CGI-specific transcription activation by the recruitment of TFIID subunits. This implies either that TFIID functions only at CGI promoters or, more plausibly, that TFIID recruitment is a key rate-limiting step of CGI promoter activation. Strong evidence attests to TFIID’s general importance at all promoters^31,32^, and rapidly depleting the central TFIID subunit Taf1 by AID (**Extended Data Fig. 6b-e, Supplementary Table 2**) indeed reduced transcription from both CGI and non-CGI promoters, and affected Hcfc1-dependent and -independent genes similarly strongly (**Fig. 4k**). This confirms that TFIID is broadly required for Pol II transcription at all promoter types and suggests that TFIID recruitment might be a key regulatory bottleneck specifically at CGI promoters.

If TFIID recruitment constitutes a critical rate-limiting step in the activation of CGI promoters, then force-recruiting TFIID should be sufficient to activate transcription at CGI promoters. To test this hypothesis, we used TetR-mediated tethering of various TFIID subunits to a CGI and a non-CGI reporter. Tethering five TFIID subunits (Taf3, Taf4b, Taf6, Taf12 and Taf13) indeed activated the CGI reporter between two- and approximately eightfold, while the non-CGI promoter was weakly activated only by Taf4b but none of the other four (**Fig. 4l**). Consistently, Taf1 is identified as a potent CGI-specific activator in the ORFtag screens above (**Extended Data Fig. 6f**). Overall, these results demonstrate that TFIID recruitment is a key rate-limiting step for transcription from CGI promoters (**Fig. 4m**).

## Discussion

Our study demonstrates that intrinsic biochemical compatibility exists between mammalian transcription co-activators and CGI promoters, the largest promoter class in mammals, suggesting there are distinct activators, rate-limiting steps, and/or mechanisms of transcription activation. We identify the first co-activators specific to CGI promoters, which activate CGI but not non-CGI promoters, even when force recruited. A founding member of this class of co-activators is Hcfc1, which has a central and essential role in mammalian gene expression and is both necessary and sufficient for the expression of thousands of genes with CGI promoters typically found at housekeeping genes that don’t respond to distal enhancer activation^33^.

Although Hcfc1 had previously been observed at CGI promoters in human cells, those studies did not establish how Hcfc1 contributes functionally to CGI promoter activation. We show that Hcfc1 directly bridges CGI-promoter-bound TFs and CoAs to the general transcription factor TFIID. Even though CoAs are broadly thought to communicate between TFs and the transcription machinery, only Mediator has been shown to act through a defined, direct TF-to-PIC bridging mechanism. We identify Hcfc1 as a second CoA with a direct bridging role and, importantly, the first to exhibit promoter-class specificity. Moreover, although TFIID is universally essential, a selective rate-limiting role for its recruitment at CGI promoters had not been recognized, nor had any CoA been shown to promote TFIID engagement in a CGI-specific manner.

The bridging of TFs to TFIID by Hcfc1 can also explain the approximate positions of transcription initiation near TF/CoA binding sites and the dispersed initiation pattern commonly observed at CGI promoters (**Fig. 4m**): CGI promoters are bound by multiple CGI-specific TFs at distinct cognate motifs. These TFs recruit Hcfc1 through their HBMs that are frequently located in intrinsically disordered regions (IDRs). Hcfc1, in turn, recruits TFIID via its AD, which is also located within a large IDR. The combination of positional variability of TF binding along the CGI promoter and structural flexibility conferred by the IDRs in both TFs and Hcfc1 allows for a range of TFIID positions and thus transcription initiation sites. Therefore, in contrast to TATA-box promoters, which display focused transcription initiation from a single nucleotide because the TATA-box–Tbp interaction positions TFIID at a defined position, the lack of a single fixed DNA motif and the involvement of flexible IDRs at CGI promoters can explain their dispersed transcription initiation patterns.

Our results further suggest that TFIID recruitment is a key regulatory bottleneck for transcription from CGI promoters, which may exist in a permissive state that renders TFIID recruitment sufficient for transcription activation without additional regulatory constraints. Despite employing a shared general transcription machinery, transcription initiation at different promoter classes may thus have distinct rate-limiting steps and be subject to distinct and exclusive compatibilities between co-activators and their promoters. This concept advances our understanding of mammalian gene regulation and suggests that other mechanisms and compatibilities for transcription regulation remain to be discovered.

## Extended data Figure legends

**Extended Data Fig. 1:**
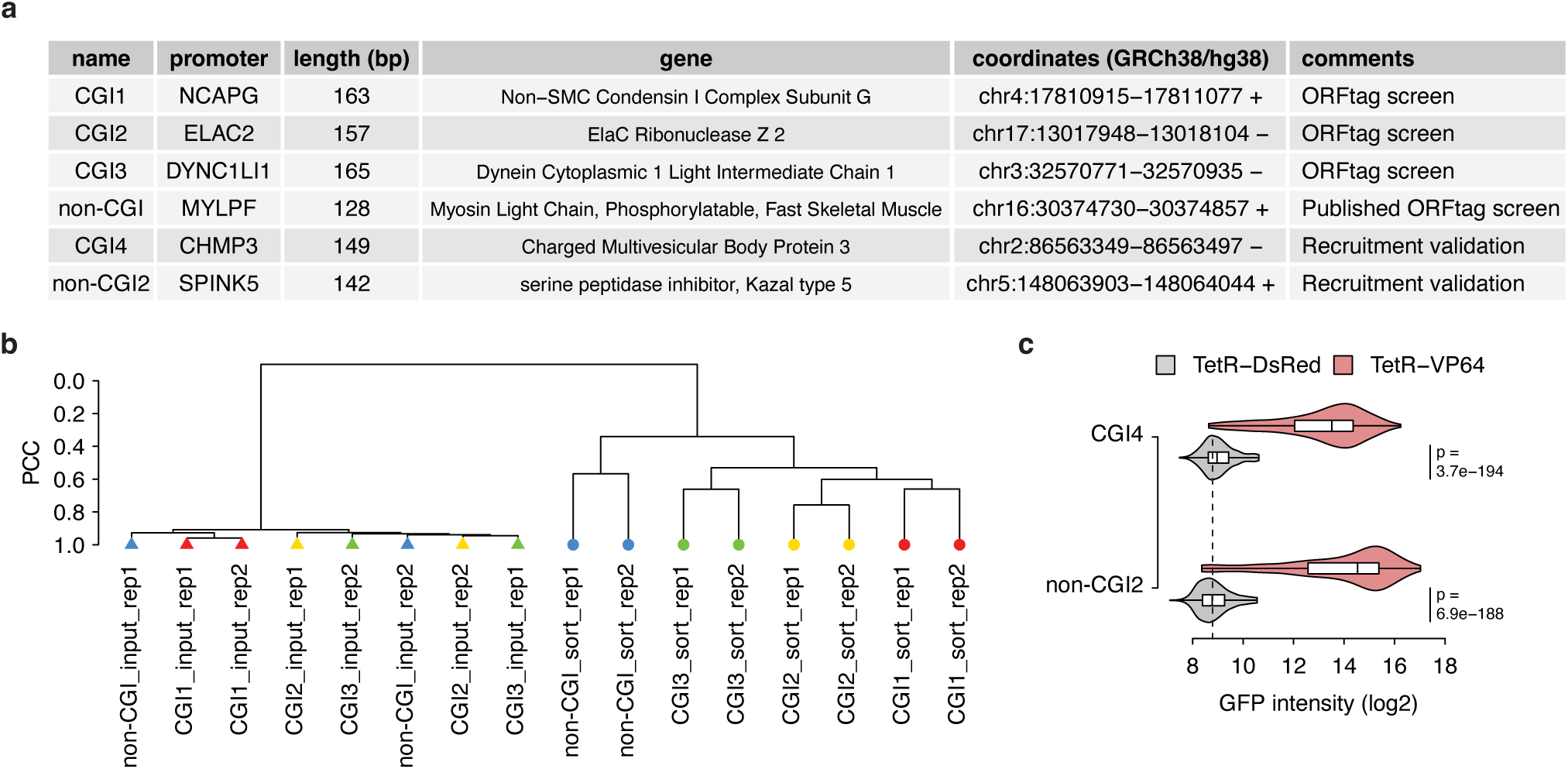
ORFtag screens and validations. **a**, Table detailing the eGFP reporter cell lines used in ORFtag screens and validations, with promoter coordinates, length, and associated gene names. **b**, Dendrogram of PCC between input control and sorted sample replicates. **c**, GFP intensity in reporter cell lines used for validations, stably expressing TetR-DsRed (control) or TetR-VP64, measured by flow cytometry. Statistical significance was assessed using a one-sided Wilcoxon test. Box plots indicate the median (central line), 25^th^-75^th^ percentiles (box), and 1.5x interquartile range (whiskers).

**Extended Data Fig. 2:**
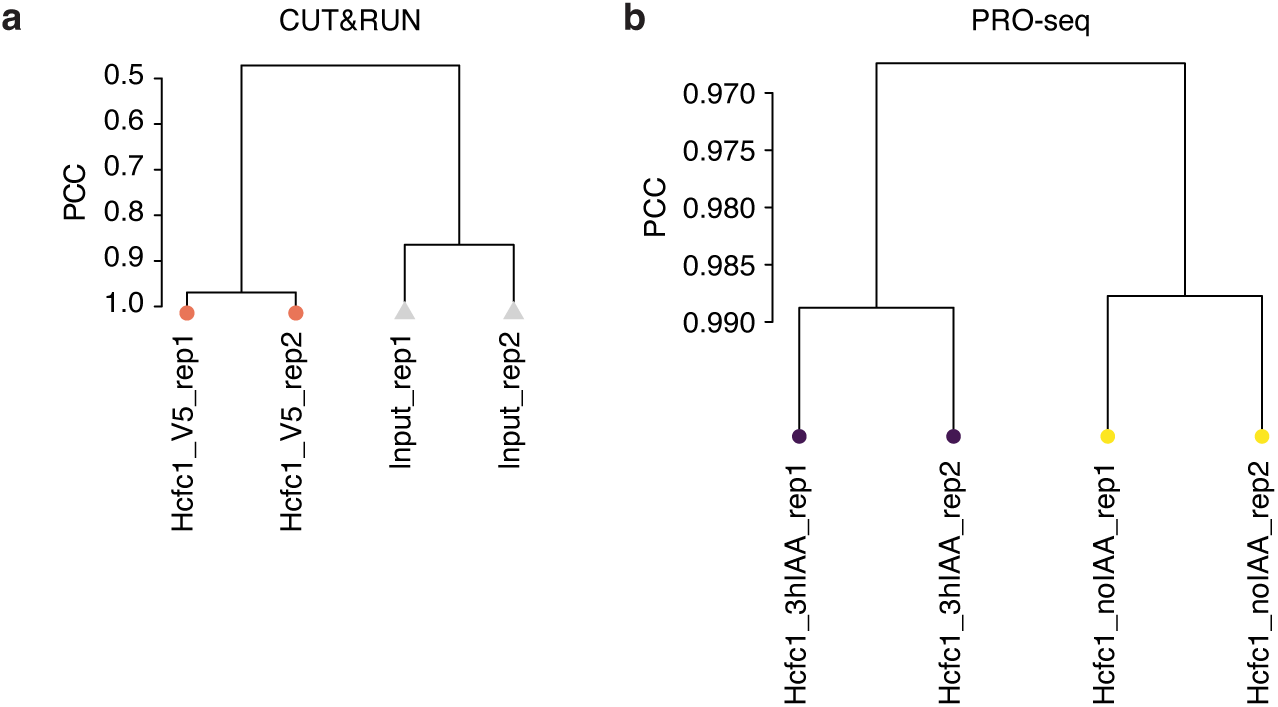
Hcfc1 PRO-seq and CUT&RUN experiments. **a**, Dendrogram of PCC comparing CUT&RUN coverage for Hcfc1 (V5 in V5-AID-Hcfc1 cell line) versus the input (V5 in parental cell line) in 10 kb bins across the genome. **b**, Dendrogram of PCC comparing spike-in normalized PRO-seq counts between biological replicates for control (no IAA) and Hcfc1-depleted (+IAA, 3 h) conditions.

**Extended Data Fig. 3:**
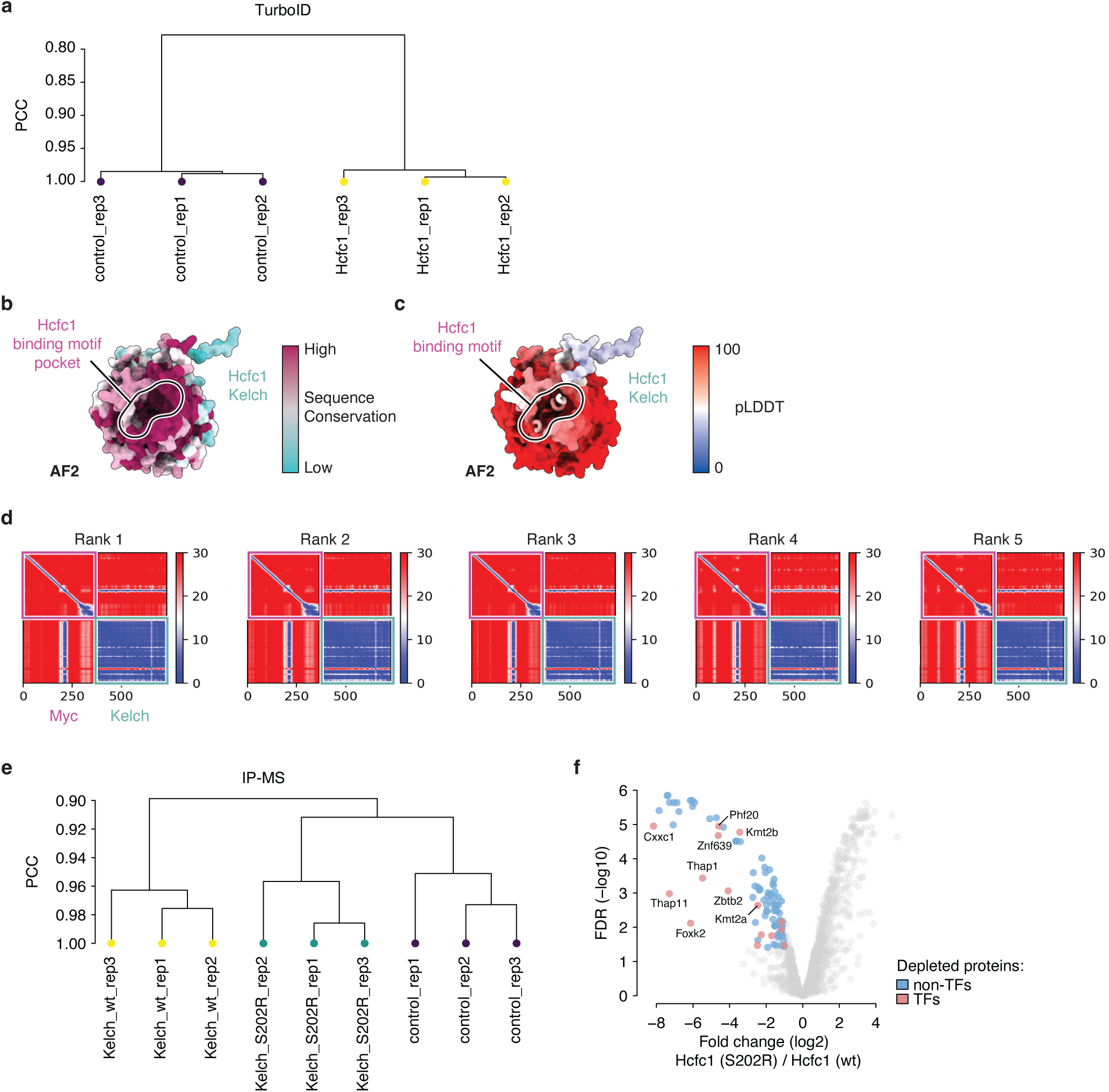
Hcfc1 TurboID and Hcfc1 Kelch domain AF2 models a, Dendrogram of PCC comparing protein abundance from TurboID mass spectrometry experiments for control and Hcfc1 baits. **b**, Sequence conservation of Hcfc1 mapped onto the surface of the AlphaFold2 (AF2)-predicted Kelch domain, highlighting the conserved HBM-binding pocket. **c**, The top-ranked AF2 model of the Hcfc1 Kelch domain complexed with its HBM-containing interactor, Myc. The surface is coloured by pLDDT (predicted local distance difference test), and only the ordered, interacting elements of the interactor are shown. **d**, The Predicted Aligned Error (PAE) plots for the AF2 model from (**c**). **e**, Dendrogram of PCC comparing protein abundance from immunoprecipitation-mass spectrometry (IP-MS) experiments for three baits: a control, the wild-type Hcfc1 Kelch domain, and the S202R mutant. **f**, Volcano plot of IP-MS results comparing wild-type Hcfc1 Kelch domain and S202R mutant interactors. Proteins significantly depleted in the mutant (log2FC > 1, FDR < 0.05) are coloured. Labelled proteins correspond to those shown in Fig. 3G.

**Extended Data Fig. 4:**
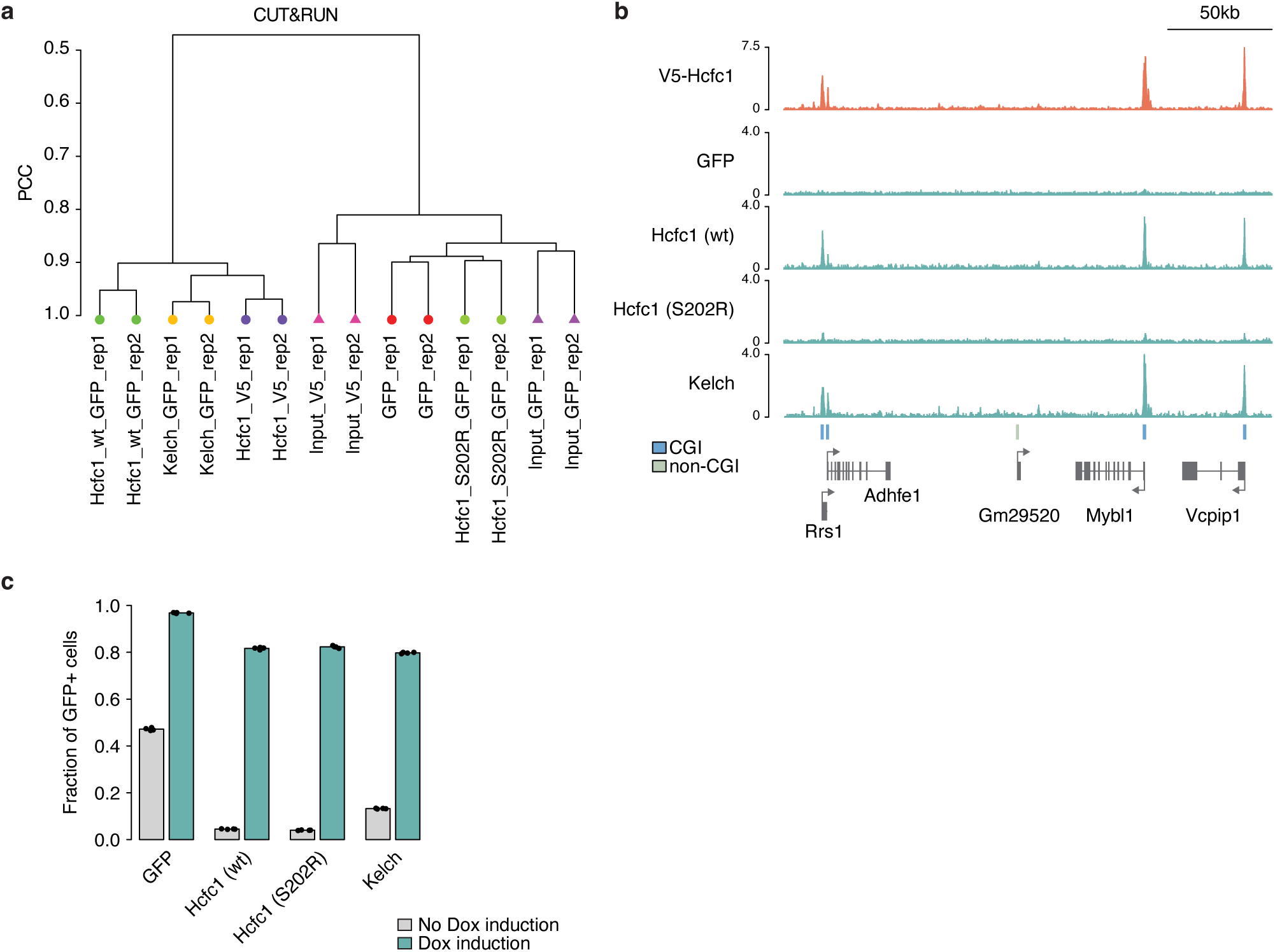
Hcfc1 rescue constructs. **a**, A dendrogram of PCC comparing CUT&RUN signal coverage across the genome in 10 kb bins. The analysis compares endogenous Hcfc1 (V5 in V5-AID-Hcfc1 cell line), its corresponding input control (V5 in parental cell line), GFP-tagged rescue constructs (GFP, after IAA treatment in V5-AID-Hcfc1 cell line), and the corresponding rescue input control (no rescue construct). **b**, Genome browser tracks comparing the Hcfc1 CUT&RUN signal between the endogenous Hcfc1 (V5) and the GFP-tagged rescue constructs. CGI (blue) and non-CGI (green) promoters are indicated for reference. **c**, Bar plot showing the fraction of cells expressing the rescue constructs before (no Doxycycline) and after induction with Doxycycline (n=4).

**Extended Data Fig. 5:**
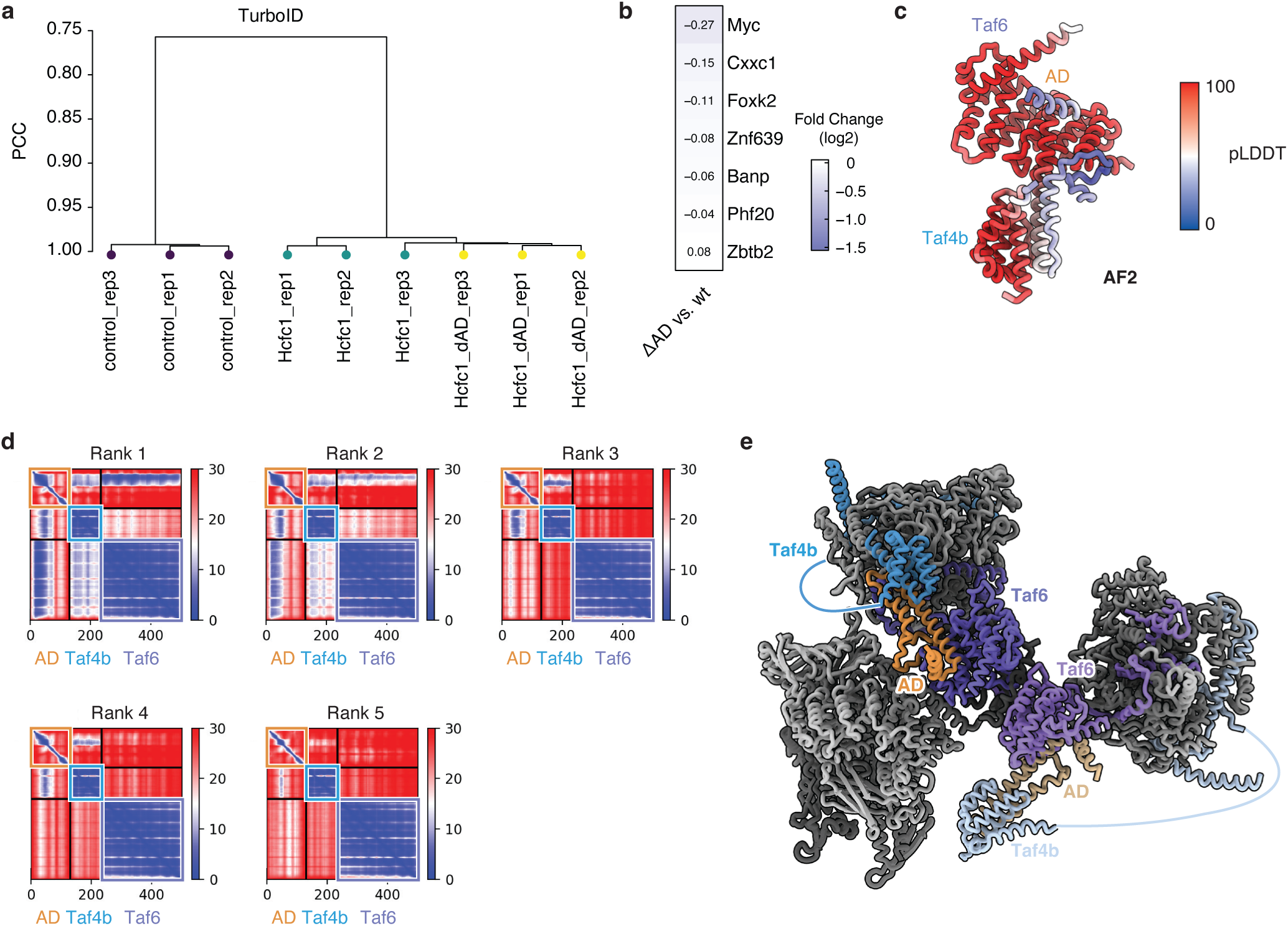
Hcfc1 activation domain interactors. **a**, Dendrogram of PCC comparing protein abundance from TurboID mass spectrometry experiments using Hcfc1 wild-type, Hcfc1 ΔAD, and control baits. **b**, Heatmap showing the relative depletion of 7 TFs in the Hcfc1 ΔAD TurboID experiment compared to wild-type, cross-referencing data from Fig. 4c and Fig. 3g. **c**, The top-ranked AF2 model of the Hcfc1-AD complexed with the interacting domains of Taf4b and Taf6. The surface is coloured by the pLDDT score. **d**, PAE plots for the AF2 models shown in (**c**). **e**, Superposition of the Hcfc1-AD-Taf4b-Taf6 AF2 model onto the experimental structure of TFIID (PDB-ID: 8GXS). The model shows that Hcfc1-AD binding is sterically compatible with the TFIID architecture, suggesting that two Hcfc1-AD copies can interact with TFIID simultaneously. TFIID is coloured in shades of grey. The primary Hcfc1-AD-Taf4b-Taf6 complex is coloured as in Fig. 4e, while the second complex is shown in lighter shades.

**Extended Data Fig. 6:**
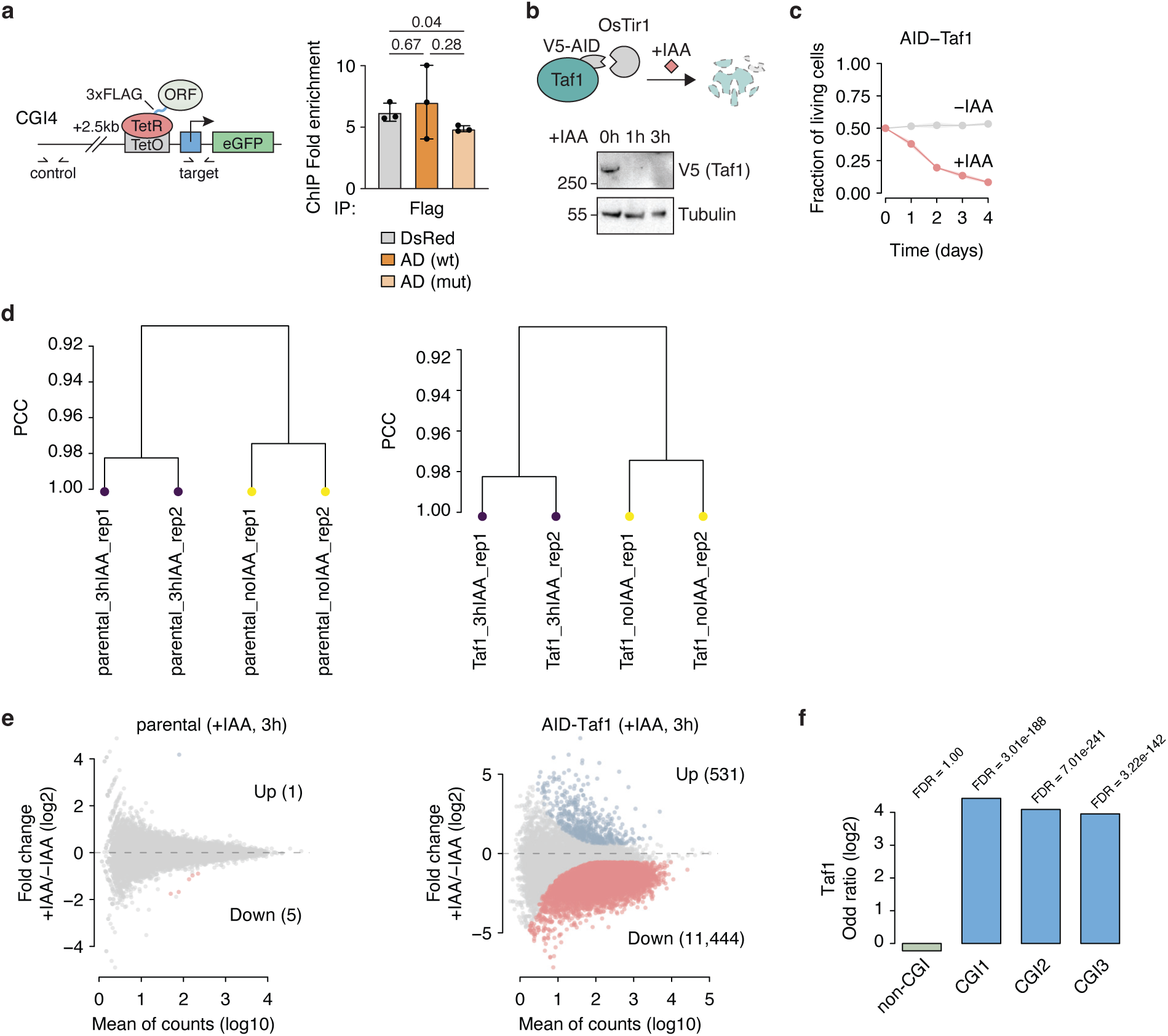
Hcfc1 activation domain and TFIID. **a**, ChIP-qPCR measuring the enrichment of FLAG-tagged TetR fusion proteins at the CGI4 reporter promoter (schematic, left). Bar graphs (right) show mean enrichment from three biological replicates, normalized to a negative control locus. Error bars represent standard deviation. Statistical significance was determined by a two-sided Student’s t-test. **b**, Schematic of the rapid Taf1 depletion using the AID system (top). The western blot (bottom) is representative of two independent experiments and shows Taf1 loss after 1 h of IAA treatment. **c**, Time-course analysis of cell viability, comparing cells with Taf1 present (-IAA, grey) to those where Taf1 is depleted (+IAA, red). Data are from two biological replicates. **d**, Dendrogram of PCC comparing spike-in normalized PRO-seq counts between biological replicates for the Taf1-depleted line and the parental control line. **e**, MA plots of PRO-seq transcriptional changes after 3 h of IAA treatment for the parental cell line (left) and AID-tagged Taf1 cell line (right). Significantly up-(blue) and down-regulated (red) genes are highlighted (FDR < 0.05, log2FC > log2(1.5)). **f**, Enrichment of Taf1 in all ORFtag screens, displayed as a bar plot of the log2 Odds Ratio (sorted vs. input). The FDR for each screen is shown above the corresponding bar.

## Supplementary Table legends

**Supplementary Data Table 1: ORFtag**

The first sheet contains the complete dataset from all ORFtag screens. For each gene, it lists the raw counts for input (count.input) and sample (count.sample) conditions, total library counts (total.input, total.sample), the log2 Odds Ratio (log2OR), and the adjusted p-value (padj). The hit column indicates significant hits. The second sheet summarizes all significant hits, specifying the number of screens in which a gene was a hit (screens_number), the names of those screens (screens_identity), and the hit classification (e.g., CGI-type or Other).

**Supplementary Data Table 2: PRO-seq**

Each sheet contains the DESeq2 results for a specific PRO-seq experiment. For each gene, it lists the base mean expression (baseMean), log2 fold change (log2FoldChange), and the adjusted p-value (padj). The diff column indicates whether a gene was significantly up-regulated, down-regulated, or unaffected.

**Supplementary Data Table 3: Proteomics**

Each sheet contains the results from a single IP-MS or TurboID experiment. For each protein identified, the table lists its UniProt accession (mouse_uniprot), gene name, and the intensity values for each replicate under the sample and control conditions. Calculated values include the log2 fold change (log2FC) and the adjusted p-value (FDR) for the sample versus control comparison. The hit column indicates significantly enriched proteins.

**Supplementary Data Table 4: Reagents and resources**

This file contains separate sheets listing the cell lines, peptides, plasmids, and oligonucleotides used in this study.

## Acknowledgements

We thank our colleagues in the Stark and Plaschka groups for their support and insightful discussions. We are grateful to Vincent Loubiere for assistance with computational analyses. We also thank the IMP/IMBA/GMI core facilities for support, particularly the Flow Cytometry and Proteomics/Mass Spectrometry teams for their outstanding service. Next-generation sequencing was performed at the Vienna Biocenter Core Facilities GmbH (VBCF) Next-Generation Sequencing Unit. We thank M. Madalinski for peptide synthesis. F.N. was supported by a Boehringer Ingelheim Fonds PhD fellowship. K.S. is supported by an EMBO postdoctoral fellowship (352-2024) and a Postdoc.Mobility fellowship from the Swiss National Science Foundation (SNSF; P500PB_225656). Research in the Plaschka group is supported by the European Research Council under the Horizon 2020 research and innovation programme (ERC-2020-STG 949081) and by the Austrian Science Fund (FWF) doc.funds program (DOC177-B). Research in the Stark group is supported by the FWF (10.55776/P29613, 10.55776/P33157, 10.55776/P36971 and 10.55776/PAT3564423) and the Vienna Science and Technology Fund (WWTF, 10.47379/LS24012). Basic research at the IMP is supported by Boehringer Ingelheim GmbH and the Austrian Research Promotion Agency (FFG, FO999902549). For the purpose of Open Access, the authors have applied a CC BY public copyright license to any Author Accepted Manuscript (AAM) version arising from this submission.

## Author contributions

F.N. generated the reporter cell lines, conducted the ORFtag screens, validated candidates, and established the AID-Hcfc1 line. S.K. generated the AID-Taf1 line. F.N. and K.S. performed the CUT&RUN experiments; F.N. and S.K. performed PRO-seq experiments. F.N., K.S., and S.K. conducted the competition assays. K.S. carried out the IP-MS and TurboID experiments, as well as the rescue assays, co-affinity purification assays, and ChIP-qPCR experiments. K.S. performed AF2 screens. F.N. performed the data processing and downstream computational analyses. M.P. assisted with experiments. C.P. and A.S. coordinated and supervised the study. All authors contributed to writing the manuscript.

## Materials and Methods

### Vectors and sequences

All vectors and sequences are listed in Table S4. Candidate sequences (ORFtag hits, Hcfc1 domains, TFIID subunits) were PCR-amplified from mES cell cDNA (Q5 High-Fidelity 2x Master Mix, NEB) or obtained from Addgene (Taf4b: 44345; Taf3: 44334; Taf6: 44337; Taf13: 44343). Inserts were cloned using Gibson cloning or Gateway technology (Invitrogen). Point mutations were introduced by round-the-horn mutagenesis^34^. All constructs were verified by Sanger sequencing or whole-plasmid sequencing (Eurofins Genomics).

### Antibodies

Primary antibodies used for Western blotting: mouse anti-V5 (Thermo Fisher, R960-25; 1:1,000), mouse anti-Hcfc1 (Novus Biologicals, NB100-68209; 1:2,000), rabbit anti-β-Tubulin (Abcam, ab6046; 1:10,000), and rabbit anti-α-Tubulin (Abcam, ab18251; 1:10,000). Secondary antibodies for Western blotting (all horseradish peroxidase (HRP)-conjugated, Cell Signaling Technology): anti-mouse (7076; 1:10,000) and anti-rabbit (7074; 1:10,000).

Antibodies used for CUT&RUN: mouse anti-V5 (Thermo Fisher, R960-25; 1 µl), rabbit anti-GFP (Thermo Fisher, A-11122; 1 µl).

Antibodies used for ChIP-qPCR: mouse IgG (Invitrogen, 31903; 3 µg), rabbit anti-Taf1 antibody (Bethyl Laboratories, A303-505A; 3 µg), mouse anti-Taf6 antibody (Invitrogen, MA3-077; 0.5 µg), rabbit anti-Med17 antibody (Abcam, ab155593; 3 µg), rabbit anti-Rpb2 antibody (Sigma Aldrich, HPA037506; 0.6 µg).

### Cell culture conditions

All experiments used diploid mouse embryonic stem (mES) cells (AN3-12) obtained from the IMBA Haplobank, Vienna, Austria. The mES cells were cultured feeder-free in high-glucose DMEM (Capricorn Scientific) supplemented with 13.5% fetal bovine serum (Sigma-Aldrich), 2 mM L-glutamine (Sigma-Aldrich), 1x penicillin-streptomycin (Sigma-Aldrich), 1x MEM nonessential amino acids (Gibco), 50 μM β-mercaptoethanol (Merck), and in-house recombinant leukemia inhibitory factor. For virus packaging, Platinum-E cells (Cell Biolabs) were maintained and transfected according to the manufacturer’s instructions. *Drosophila* S2 cells (Thermo Fisher) were cultured in Schneider’s Drosophila Medium (Gibco) containing 10% heat-inactivated fetal bovine serum (Sigma-Aldrich). Mammalian cells were maintained at 37 °C and 5% CO2, whereas S2 cells were kept at 27 °C and 0.4% CO2. All cell lines were routinely tested for mycoplasma contamination.

### Reporter cell lines

The parental cell line was established previously^35^ and contains a BFP-expressing cassette integrated at a stably expressing locus on chromosome 15, compatible with Flp recombinase-mediated cassette exchange (RMCE). To generate new reporter lines, 5 million cells were electroporated using a Maxcyte STX device (GOC-1, Opt5 program) with 10 µg FRT/F3-flanked reporter plasmid and 6 µg Flp recombinase-expressing plasmid in MaxCyte electroporation buffer. 7 days post-transfection, cells were sorted by FACS for BFP-negative cells (BD FACSAria III or IIu), and clonal populations were established. Genotyping of clones was performed by integration site-specific PCR and confirmed by Sanger sequencing. The reporter construct consists of a PuroR-IRES-eGFP cassette driven by a minimal promoter (see Extended Fig. 1A), with 7 TetO sites flanked by loxP sequences upstream of the promoter. Final clones were tested for low basal expression and robust inducibility using TetR-DsRed and TetR-VP64 constructs, respectively.

### ORFtag screens

A mixture of 3 retroviral ORFtag plasmids (Addgene IDs: 220981, 220982, 220983) at a 1:1:1 mass ratio was used for ORFtag screens, as previously described^14^. Each plasmid contains all elements required for the inverse-PCR protocol, including TetR with an N-terminal nuclear localization signal, a 2x GGGS linker, BC2-tag, and 3x FLAG-tag, a consensus splice donor motif, and a segment of the Hprt intron. To allow tagging in all 3 reading frames, the plasmids were designed with the insertion of 0, +1, or +2 nucleotides immediately before the splice donor site (GT). This results in the following sequences: AAG-CAG-GT for frame 1, AAG-G-CAG-GT for frame 2, and AAG-GC-CAG-GT for frame 3, where AAG corresponds to the last codon of the 3x FLAG tag.

For virus production, the ORFtag plasmid mix was packaged in Platinum-E cells using polyethylenimine (PEI) transfection. Platinum-E cells (11.25 million per 150 mm dish) were seeded 24 h prior to transfection. The transfection mix, prepared in high-glucose DMEM (no supplements), contained 45 µg ORFtag plasmid mix, 15 µg pCMV-Gag-Pol (Cell Biolabs), and 135 µl PEI in a final volume of 3.2 ml per dish. After 20 min incubation, the mixture was added to the cells. Medium was replaced with mES medium after 12 h. Viral supernatants were collected at 24 and 40 h post-transfection, pooled, filtered (0.45 µm filter), and supplemented with 6 µg/ml polybrene (Sigma).

Reporter cell lines (150 million) were seeded in 245x245 mm dishes 4 h before transduction and infected with the retroviral supernatant, targeting < 15% transduction efficiency to ensure single-copy integration. After 24 h, cells were re-seeded in medium with 0.1 mg/ml G418 (Gibco) for selection, which continued until no viable cells remained in the non-transduced control dish (typically 4–5 days), with daily PBS washes and medium changes. After selection, 40 million cells were harvested as the unsorted input for integration site mapping. The remaining cells were sorted by FACS for GFP-positive cells (BD FACSAria III or IIu) and processed for integration site mapping.

Transduction efficiency was determined for each screen by plating 10,000 cells per 150 mm dish under G418 selection, alongside a control dish of 1,000 cells without selection. After 10 days, colony counts were used to calculate transduction efficiency, which ranged from 2.6–12.1%.

### Inverse-PCR followed by next-generation sequencing

Genomic locations of ORFtag integrations were identified using a modified inverse-PCR coupled with next-generation sequencing (iPCR-NGS) protocol, as previously described ^14^. In brief, genomic DNA was isolated by lysing cell pellets overnight at 55 °C in lysis buffer (10 mM Tris-HCl pH 8.0, 5 mM EDTA, 100 mM NaCl, 1% SDS, 0.5 mg/ml proteinase K). After lysis, samples were treated with RNase A (Qiagen, 100 mg/ml, 1:1,000 dilution) for 2 h at 37 °C, followed by 2 rounds of phenol:chloroform:isoamyl alcohol extraction and a final extraction with chloroform:isoamyl alcohol. Up to 4 µg of genomic DNA was digested overnight in separate reactions with NlaIII and MseI (NEB) at 37 °C, then purified using the Monarch PCR & DNA Cleanup Kit (NEB). Ring ligation was performed overnight at 16 °C with T4 DNA ligase (NEB), followed by heat inactivation (65 °C, 15 min) and linearization with SbfI-HF (NEB) for 2 h at 37 °C. The DNA was again purified (Monarch PCR & DNA Cleanup Kit) and amplified by nested PCR using KAPA HiFi HotStart ReadyMix (Roche) and specific primers (FN865, FN866) for 16 cycles. After purification with AMPure XP Reagent (Beckman Coulter, 1:1 beads:PCR product), a second round of iPCR amplification was performed for 18 cycles using KAPA HiFi HotStart ReadyMix and a second primer pair (FNR848, UD7xx containing an 8-nt barcode). Amplified libraries were size-selected to 400–800 bp using SPRIselect beads (Beckman Coulter). Sequencing was performed on an Illumina NextSeq550 platform using a custom first-read primer (1:1 mix of FN860 and FN861). A complete list of primers is provided in Table S4.

### Individual recruitment validations

Candidate sequences were cloned into a retroviral destination vector containing a PGK promoter, puromycin resistance (PuroR), and an N-terminal tag (TetR with NLS, 2x GGGS linker, BC2 tag, 3x FLAG). The TetR-candidate and PuroR coding sequences are separated by an IRES. All primers and plasmids are listed in **Supplementary Table 4.**

Retroviral constructs were packaged in Platinum-E cells (see above). 0.8 million cells per well were seeded in 6-well dishes 24 h prior to transfection. Transfection mix (in 215 µl high-glucose DMEM): 3.2 µg TetR-candidate plasmid, 1 µg pCMV-Gag-Pol and 9.6 µl PEI was incubated for 20 min, then added to cells. Medium was changed to mES medium after 12 h. Viral supernatant was collected 24 h post-transfection, filtered (0.45 µm), and supplemented with 6 µg/ml polybrene (Sigma). Reporter cell lines (170,000 cells per well in a 24-well plate) were transduced, harvested 24 h later, and plated in medium containing 1 µg/ml puromycin (InvivoGen) for selection. After 5 days of selection, reporter expression was analyzed using an LSR Fortessa (BD) flow cytometer. Data were processed and visualized using FlowJo (v10.10) and R/flowCore (v2.12.2).

### AID cell line generation

A parental cell line stably expressing the Oryza sativa Tir1 (OsTir1) E3 ligase for the AID system was previously established^14^. To generate N-terminally AID-tagged Hcfc1 and Taf1 cell lines, 5 million parental cells were transfected with 10 μg of a plasmid encoding Cas9 and a guide RNA targeting either the Hcfc1 (TGCATCCTTAACCGGAACTA) or Taf1 (TATTTTTTCTTTCCCGCCCA) locus, along with 5 μg of a donor plasmid containing a Blasticidin-P2A-V5-AID-GGGS knock-in cassette flanked by 20 bp microhomology arms to facilitate knock-in via microhomology-mediated end joining^36^. Transfection was performed using a Maxcyte STX device (GOC-1, Opt5 program). 2 days post-transfection, selection was performed using 10 μg/ml Blasticidin (Thermo Fisher). Individual clones were genotyped by knock-in-site-specific PCR and Sanger sequencing (see Table S4 for primer sequences) to identify homozygous clones. Candidate clones were further validated by western blotting for the integrated V5 tag. Auxin-dependent protein depletion was confirmed following a 1 h and 3 h treatment with 500 μM indole-3-acetic acid (IAA; Merck).

### PRO-seq

PRO-seq was performed following a parallelized protocol adapted from^37^; with 2 biological replicates for each condition: cells treated for 3 h with 500 µM IAA (Merck) and untreated control. For each sample, 20 million mES cells were harvested (5 min, 500 *g*, 4 °C), washed with ice-cold PBS. Spike-in control (S2 *Drosophila* cells; 1% of mES cells) was added during the following nuclei permeabilization step. Cells were resuspended in 20 ml ice-cold permeabilization buffer (10 mM Tris-HCl pH 7.5, 300 mM Sucrose, 10 mM CaCl2, 5 mM MgCl2, 1 mM EGTA, 0.05% Tween-20, 0.1% NP-40, 0.5 mM DTT, and protease inhibitors), incubated on ice for 5 min, and the nuclei were pelleted (5 min, 1,000 *g*, 4 °C), washed 2 times, and then resuspended in 200 µl storage buffer (10 mM Tris-HCl pH 8.0, 25% Glycerol, 5 mM MgCl2, 0.1 mM EDTA, 5 mM DTT). The nuclear run-on (NRO) was initiated by mixing 10 million nuclei (100 µl) with 100 µl of 2x NRO mix (10 mM Tris-HCl pH 8.0, 5 mM MgCl₂, 300 mM KCl, 1 mM DTT, 1% Sarkosyl, 0.25 mM ATP/GTP, 0.05 mM Biotin-11-CTP/UTP, 0.8 U/µl Murine RNase Inhibitor, and incubated for 3 min at 37 °C. The reaction was stopped with 500 µl Trizol LS (Invitrogen). RNA was extracted using chloroform separation and ethanol precipitation, then resuspended in 50 µl of water. The RNA was subsequently hydrolyzed with 5 µl of 1 M NaOH on ice for 15 min, neutralized with 25 µl of 1 M Tris-HCl pH 6.8, and purified using a Bio-Rad P-30 column. Biotinylated RNA was enriched using 50 µl Streptavidin M-280 Dynabeads (20 min, room temperature; Thermo Fisher), then washed 2 times with high-salt buffer (50 mM Tris-HCl pH 7.4, 2 M NaCl, 0.5% Triton X-100), 2 times with binding buffer (10 mM Tris-HCl pH 7.4, 300 mM NaCl, 0.1% Triton X-100), and once with low-salt buffer (5 mM Tris-HCl pH 7.4, 0.1% Triton X-100). RNA was extracted with Trizol, purified with Zymo Direct-zol kit, and eluted into 5 µl 3’ RNA adapter (Rev3, 1:8 dilution, 10-nt UMI) solution. 3’ adapter ligation was performed overnight at 16 °C with T4 RNA Ligase I (50% PEG), then RNA was rebound to streptavidin beads. All subsequent enzymatic steps were performed on-bead: 5’ cap was removed with TAP (1 h, 37 °C), then 5’ end phosphorylated with T4 PNK (1 h, 37 °C). 5’ RNA adapter (Rev5) was ligated (4 h, room temperature, T4 RNA Ligase I), then RNA was washed, eluted with Trizol, and purified with Zymo Direct-zol, eluting in 10.5 µl water. RNA library was reverse transcribed with SuperScript III (RP1 primer), and the resulting cDNA was amplified with KAPA HiFi HotStart kit (RP1 reverse primer, RPI index forward primer), monitoring by qPCR and stopping in exponential phase. PCR products were purified with 1.4x AMPure XP beads, quantified, and sequenced on an Illumina NextSeq550.

### PiggyBac cell line generation

To generate mES cells carrying a Doxycycline (Dox)-inducible target gene, a PiggyBac transposon system was used^38^. Expression constructs were introduced downstream of a 3xFlag-NLS-GFP or 3xFlag-NLS-TurboID tag under the control of Dox-inducible promoter. In addition, the integration cassette also contained the tTA activator and the puromycin resistance gene under the control of a PGK promoter.

10 million cells were electroporated using a Maxcyte STX device (GOC-1, Opt5 program) with 7.5 µg plasmid encoding the PiggyBac transposase and 7.5 µg of the expression construct plasmid in MaxCyte electroporation buffer (see **Supplementary Table 4** for details). 1 day post-transfection, selection was performed using 1 μg/ml Puromycin (InvivoGen). After 5 days of continuous selection, the stable polyclonal cell line was used for downstream experiments.

### CUT&RUN

CUT&RUN was performed with 2 replicates of 1 million cells each, following the EpiCypher CUTANA v2.1 protocol with modifications^39^; all steps were performed in 8-strip PCR tubes. For the experiment performed under depletion-rescue conditions, endogenously V5-AID-tagged Hcfc1 mES cells carrying Dox-inducible rescue constructs were used. Cells were seeded in duplicate for each condition and cultured for two days. After 24 hours, rescue construct expression was induced by adding Dox (0.05 µg/ml) to the medium. To rapidly deplete endogenous Hcfc1, IAA (Merck) was added to a final concentration of 500 µM three hours prior to harvesting.

For each sample, 1 million cells were harvested (600 *g*, 3 min) and washed 2 times in 100 μl Wash Buffer (20 mM HEPES-NaOH pH 7.5, 150 mM NaCl, 0.5 mM Spermidine) supplemented with 1x cOmplete EDTA-free protease inhibitor cocktail (Roche). The washed cells were then resuspended in 100 μl Wash Buffer. Concanavalin A (ConA) conjugated beads (EpiCypher; 10.5 μl/reaction) were activated by 2 washes in 100 μl cold Bead Activation Buffer (20 mM HEPES-KOH pH 7.9, 10 mM KCl, 1 mM CaCl2, 1 mM MnCl2), then resuspended in 10.5 μl of the same buffer. Washed cells were mixed with 10 μl activated ConA beads, and incubated for 10 min at room temperature. Cell-bead complexes were collected on a magnetic rack, resuspended in 100 μl cold Antibody Buffer (Wash Buffer + 0.0025% Digitonin + 2 mM EDTA), and incubated overnight at 4 °C with 1 μl antibody (mouse anti-V5 [Thermo Fisher, R960-25] or rabbit anti-GFP [Thermo Fisher, A-11122]). The next day, samples were collected on a magnetic rack, washed 2 times with 200 μl cold Digitonin Buffer (Wash Buffer + 0.0025% Digitonin), resuspended in 150 μl Digitonin Buffer, and incubated for 10 min at room temperature with 3.5 μl pAG-MNase (30 µg/ml). The samples were then washed 2 times with 200 μl cold Digitonin Buffer and resuspended in 50 μl of the same buffer. For digestion, tubes were chilled on ice, 1 μl 100 mM CaCl2 was added, and samples were incubated for 2 h at 4 °C on a nutator. Digestion was stopped with 33 μl STOP Buffer (340 mM NaCl, 20 mM EDTA, 4 mM EGTA, 50 µg/ml RNase A, 50 µg/ml Glycogen). Samples were incubated 10 min at 37 °C to release DNA. After pelleting the beads on a magnetic rack, the supernatant was collected. DNA was purified with NEB Monarch PCR & DNA Cleanup Kit, eluted in 12 μl Elution Buffer. Purified DNA was used immediately for library preparation using the NEBNext Ultra II DNA Library Prep Kit (no size selection) and sequenced on an Illumina NextSeq550.

### TurboID

For the TurboID proximity labelling experiment, mES cells were utilized that expressed one of three Dox-inducible constructs: Hcfc1 wild-type (wt), Hcfc1 ΔAD, or a TurboID tag alone (negative control). Cells for each condition were seeded in triplicate and cultured for 2 days. After the first 24 h, transgene expression was induced by the addition of 0.05 µg/ml Dox to the medium. 3 h prior to harvesting, endogenously V5-AID tagged Hcfc1 was degraded by adding IAA to a final concentration of 500 µM. Proximity labelling was then initiated by adding Biotin to a final concentration of 500 µM, followed by a 10-minute incubation at 37 °C.

Roughly 10 million cells were harvested and washed once with 1x PBS. Nuclei were isolated by resuspending cells in swelling buffer (10 mM Tris-HCl pH 7.5, 2 mM MgCl2, 3 mM CaCl2) supplemented with 1x cOmplete EDTA-free protease inhibitor cocktail (Roche), followed by incubation on ice for 10 min. Cells were pelleted (5 min, 500 *g*, 4 °C), the supernatant was removed, and the pellets were resuspended in GRO lysis buffer (10 mM Tris-HCl pH 7.5, 2 mM MgCl2, 3 mM CaCl2, 0.5% NP-40, 10% glycerol) supplemented with 1x cOmplete EDTA-free protease inhibitor cocktail (Roche). The suspension was incubated at 4 °C with rotation for 30 min. Nuclei were collected (5 min, 700 *g*, 4 °C) and lysed in RIPA buffer (50 mM Tris-HCl pH 7.5, 150 mM NaCl, 0.1% SDS, 0.5% Sodium Deoxycholate, 1% Triton X-100, 2 mM DTT) supplemented with 1x cOmplete EDTA-free protease inhibitor cocktail (Roche) and Benzonase (50 U/ml). The lysate was incubated for 1 h at 4 °C, and insoluble parts were removed by centrifugation (10 min, 18,000 g, 4 °C).

In parallel, 50 μl beads/sample (Dynabeads™ M-280 Streptavidin; Thermo Fisher) were equilibrated in RIPA buffer. Beads were added to the cleared lysate, incubated at 4 °C overnight with rotation. The following day, beads were collected on a magnetic rack and washed sequentially with: RIPA buffer, Wash buffer 1 (2% SDS), Wash buffer 2 (50 mM HEPES pH 7.5, 500 mM NaCl, 1 mM EDTA, 0.1% Sodium Deoxycholate, 1% Triton X-

100), Wash buffer 3 (10 mM Tris-HCl pH 7.5, 250 mM LiCl, 1 mM EDTA, 0.5% Sodium Deoxycholate, 1% NP-40) and 4 washes with non-detergent washing buffer (20 mM Tris-HCl pH 7.5, 100 mM NaCl). After the final wash, all buffer was removed, and the beads were snap-frozen in liquid nitrogen. Bound proteins were analyzed by mass spectrometry starting from an on-bead digest.

### Immunoprecipitations followed by mass spectrometry

For immunoprecipitation followed by mass spectrometry (IP-MS), mES cells with Dox-inducible expression of specific constructs were utilized. The constructs included Hcfc1-Kelch wt, Hcfc1-Kelch S202R, and a GFP negative control, each bearing an N-terminal TetR-3xFLAG-NLS tag. Cells were seeded and cultured for 2 days. After 1 day, transgene expression was induced by replacing the medium with one containing 3 µg/ml Dox. 3 h prior to harvesting, IAA (Merck) was added to a final concentration of 500 µM to induce the degradation of endogenously tagged Hcfc1.

Roughly 250 million cells were harvested and washed once with 1x PBS. Nuclear extract was generated as described above. Nuclei were resuspended in NET buffer (50 mM Tris-HCl pH 7.5, 150 mM NaCl, 0.1% Triton X-100, 10% glycerol) supplemented with 1x cOmplete EDTA-free protease inhibitor cocktail (Roche) and Benzonase (50 U/ml) and lysed by sonication (45 s, 40% output). Insoluble fragments were precipitated by centrifugation (15 min, 20,000 g, 4 °C). Supernatants were incubated with Anti-FLAG M2 Magnetic Beads (Merck), pre-equilibrated in NET buffer, overnight at 4 °C with rotation. Samples were washed 4 times with NET buffer and 4 times with NET buffer lacking detergents and protease inhibitors (50 mM Tris-HCl pH 7.5, 150 mM NaCl, 10% glycerol). After the final wash, the buffer was removed, and the beads were snap-frozen in liquid nitrogen. Bound proteins were analyzed by mass spectrometry starting from an on-bead digest.

### ChIP-qPCR

100 million mES cells stably expressing TetR fusions with DsRed, AD or AD mutant under the control of a PGK promoter (see *Individual recruitment validations*) were harvested and washed once with 1x PBS. Cells were resuspended in 1x PBS and crosslinked for 15 min with 1.1% (v/v) formaldehyde (Sigma) while rotating at room temperature. The reaction was quenched by adding glycine to a final concentration of 125 mM followed by incubation for 5 min at 4 °C. Cells were collected by centrifugation (5 min, 1,200 g, 4 °C).

For nuclei preparation, cell pellets were resuspended in CiA NP-Rinse 1 buffer (50 mM HEPES pH 8.0, 140 mM NaCl, 1 mM EDTA, 10% glycerol, 0.5% NP-40, 0.25% Triton X-100) supplemented with 1x cOmplete EDTA-free protease inhibitor cocktail (Roche) and incubated on ice for 10 min. Nuclei were collected by centrifugation (5 min, 1,200 g, 4 °C), resuspended in CiA NP-Rinse 2 buffer (10 mM Tris pH 8.0, 200 mM NaCl, 1 mM EDTA, 0.5 mM EGTA) supplemented with 1x cOmplete EDTA-free protease inhibitor cocktail (Roche). Nuclei were collected again by centrifugation (5 min, 1,200 g, 4 °C) and the resulting pellet was washed gently 2 times with CiA Covaris Shearing Buffer (10 mM Tris-HCl pH 8.0, 1 mM EDTA, 0.1% SDS) supplemented with 1x cOmplete EDTA-free protease inhibitor cocktail (Roche). Finally, the nuclei were resuspended in 2.7 ml ice-cold CiA Covaris Shearing Buffer. Chromatin was sheared using a Covaris E220 (duty factor 5%, PiP 140 W, Cycles/burst (CPB) 200) operating at 4 °C. Insoluble debris was removed by centrifugation (15 min, 20,000 g, 4 °C).

To generate input samples, 1 % of total chromatin was used. Input samples were first treated with RNase A (Thermo Fisher) for 30 min at 37 °C, then digested with Proteinase K (in-house) for 2.5 h at 55 °C, followed by overnight incubation at 65 °C to reverse crosslinking. Sheared chromatin was extracted using the phenol:chloroform:isoamylalcohol method followed by isopropanol precipitation. The resulting pellet was resuspended in 25 µl 1xTE (10 mM Tris-HCl pH 8.0, 0.1 mM EDTA). Shearing efficiency was assessed by agarose gel electrophoresis.

For pre-clearing, chromatin was incubated with control agarose beads (Pierce) for 1 h at 4 °C with overhead shaking. For the immunoprecipitation reactions, 150 μg chromatin in 500 µl 1x IP buffer (50 mM HEPES pH 7.5, 150 mM NaCl, 1 mM EDTA, 1% Triton X-100, 0.1% DOC, 0.1% SDS) was incubated with the respective antibodies (3 µg mouse IgG [Invitrogen, 31903]; 3 µg anti-Taf1 antibody [Bethyl Laboratories, A303-505A]; 0.5 µg anti-Taf6 antibody [Invitrogen, MA3-077]; 3 µg anti-Med17 antibody [Abcam, ab155593]; 0.6 µg anti-Rpb2 antibody [Sigma Aldrich, HPA037506]) overnight at 4 °C with overhead shaking. In parallel, 20 µl protein G dynabeads (Thermo Fisher) per sample were washed 3 times with blocking solution (1 mg/ml BSA in 1x PBS) and incubated overnight at 4 °C with rotation.

The next day, the blocked beads were collected on a magnetic rack and washed 5 times with 1x IP buffer. 20 µl beads were added to each sample and incubated for at least 6 h at 4 °C with rotation. Beads were collected on a magnetic rack and washed 5 times with 1x IP buffer, 3 times with DOC buffer (10 mM Tris pH 8.0, 250 mM LiCl, 1 mM EDTA, 0.5% NP-40, 0.5% sodium deoxycholate) and once with 1x TE containing 50 mM NaCl. Chromatin was eluted from the beads in two steps using a total volume of 300 µl elution buffer (100 mM NaHCO3, 1% SDS), incubated at 65 °C for 20 min each. Eluates were treated with RNase A (Thermo Fisher) for 30 min at 37 °C, then digested with Proteinase K (in-house) at 55 °C for 2.5 h, then incubated overnight at 65 °C to reverse crosslinking. DNA fragments were extracted using phenol:chloroform:isoamylalcohol method followed by isopropanol precipitation. Pellets were resuspended in 50 µl 1x TE.

Each qPCR reaction was assembled in a 20 µl volume, consisting of 10 µl of 2x GoTaq qPCR Master Mix (Promega), 1 µl of the respective primer set (10 µM; see **Supplementary Table 4**), 1 µl of immunoprecipitated chromatin or diluted input chromatin (1:10), and 8 µl of nuclease-free water. Relative enrichment was calculated using the 2^-ΔΔCT^ method, first normalizing to input, then to a negative control region located 2.5 kb upstream of the TSS.

### Protein expression in E. coli

The following constructs were expressed in *E. coli* BL21 Star (DE3) cells (Invitrogen): pOPINB+-10His-3xFLAG-MmTaf4b250-353-3C-MmTaf6219-484 or pOPINB+-10His-3xFLAG-

MmTaf4b250-353-L310R-3C-MmTaf6219-484-I280R. Cells were grown at 37 °C in LB medium with appropriate antibiotics until OD600 = 0.5 was reached. Protein expression was induced by adding 1 mM isopropyl-β-D-thiogalactopyranoside (IPTG), followed by overnight incubation at 20 °C. Cells were harvested by centrifugation (15 min, 4,000 g, 4 °C).

### Purification of proteins

To purify MmTaf4b-MmTaf6 fusion constructs (wt and LI2RR mutant), cell pellets were resuspended in lysis buffer 1 (25 mM HEPES-NaOH pH 7.9, 500 mM NaCl, 5% glycerol, 20 mM imidazole, 2 mM β-mercaptoethanol) supplemented with 1x cOmplete EDTA-free protease inhibitor cocktail (Roche), 1 mg/ml lysozyme and 5 μg/ml DNase I. Cells were lysed by sonication (60% amplitude, total time 2:30 min, 1 s on / 1 s off pulses). The cleared lysate was filtered (0.45 μm) and loaded onto HisTrap columns (Cytiva).

The columns were washed sequentially with lysis buffer 1 and then lysis buffer 1 containing 50 mM imidazole. Bound proteins were eluted using a linear gradient up to 300 mM imidazole in lysis buffer 1. The eluate was diluted to 100 mM salt and applied to a HiTrapQ anion exchange column (Cytiva). Proteins were eluted by a linear gradient to 500 mM salt.

Finally, size-exclusion chromatography was performed using a Superdex75 column (Cytiva) pre-equilibrated in gel filtration buffer (25 mM HEPES pH 7.9, 250 mM NaCl, 5% glycerol, 1 mM DTT). Peak fractions containing the target protein were concentrated and snap-frozen in liquid nitrogen.

### Co-affinity purification assays

For co-affinity purification (co-AP) assays, synthetic activation domain peptides (wt or mutant), which were N-terminally biotinylated and C-terminally coupled to a fluorescein moiety, were first immobilized by binding them in excess to 25 µl of Dynabeads M-280 Streptavidin (Thermo Fisher) per sample. Binding was performed in pull-down buffer (10 mM HEPES pH 7.5, 50 mM NaCl, 5% glycerol, 0.01% NP-40, 2 mM DTT) supplemented with protease inhibitors for 45 min at 4 °C with rotation. Beads were collected on a magnetic rack and washed 3 times with pull-down buffer. Recombinant MmTaf4b-MmTaf6 fusions (wt or mutant) were added to the respective samples at a final concentration of 20 μM. The samples were incubated for 1 h at 4 °C with rotation. After the incubation, the beads were collected on a magnetic rack and washed 3 times with pull-down buffer to remove unbound proteins. The bound protein complexes were then eluted from the beads using protein sample buffer. Proteins were separated by SDS-PAGE. Loading of synthetic peptides was detected by in-gel fluorescein fluorescence, while interacting proteins were detected by Coomassie staining.

### Cell viability time course

To evaluate the effect of AID-mediated depletion of tagged proteins on cell viability, 0.1 million cells of tagged cell lines and wild-type mES cells were seeded in a 1:1 ratio with or without 500 µM IAA (Merck). The ratio of endogenously tagged cells and wild-type cells was determined using Flow Cytometry (iQue Screener Plus, Sartorius) based on the mCherry expressed in cell lines containing the OsTir enzyme^14^.

For rescue experiments, the mES cells carrying Dox-inducible rescue constructs were pre-cultured for 24 hours in either standard medium (uninduced) or medium supplemented with 0.05 µg/ml Dox (induced). 0.1 million cells of induced or uninduced cells and wild-type mES cells were seeded in a 1:1 ratio with or without 500 µM IAA (Merck). Ratio of tagged cell lines and wild-type cells was determined using Flow Cytometry (iQue Screener Plus, Sartorius) based on the mCherry expressed in cell lines containing the OsTir enzyme. In addition, GFP levels were monitored reflecting expression levels of the rescue constructs. Data were processed and visualized using FlowJo (v10.10) and R/flowCore (v2.12.2).

### Western blotting

3 million mES cells were washed with 1x PBS and were lysed directly in 300 μl 2x protein sample buffer. The lysates were denatured at 95 °C for 5 min and to solubilize chromatin, samples were sonicated by a Diagenode Bioruptor Pico (Diagenode) for 2 min (30 s sonication followed by 30 s breaks) operating at 4 °C. Proteins were separated by SDS-PAGE, transferred onto PVDF membrane and blocked with 5 % milk powder in 1x TBST followed by an overnight incubation with the primary antibody at 4 °C. Membranes were washed 3 times with 1x TBST for 5 min and incubated with appropriate HRP conjugated secondary antibody for 1 h. Membranes were washed 3 times with 1x TBST for 5 min and were visualized using Clarity ECL substrate (Bio-Rad) and a ChemiDoc (Bio-Rad).

### Bioinformatics analyses

All bioinformatic analyses were performed in R (v4.3.0, https://www.R-project.org/). Genomic coordinate computations were performed using the GenomicRanges (v1.54.1,^40^) and data.table (v1.17.0, https://CRAN.R-project.org/package=data.table) packages. All box plots indicate the median (central line), 25th-75th percentiles (box), and 1.5x interquartile range (whiskers); outliers are omitted.

### ORFtag analysis

iPCR sequencing reads were trimmed to remove Illumina adapter sequences using Trim Galore (v0.6.0) with default parameters. The trimmed reads were aligned to the mm10 mouse genome assembly using Bowtie2 (v2.3.4.2,^41^) with default settings; for paired-end samples, only the first mate was considered. Aligned reads were filtered to remove PCR duplicates and reads with mapping quality scores ≤ 30 using samtools (v1.9,^42^). Each mapped insertion was then assigned to the nearest downstream exon junction within a maximum distance of 200 kb, based on GENCODE vM25 annotations. Only exons from protein-coding transcripts were considered, excluding the first exon of each transcript (as it lacks splicing acceptor sites). Intronless transcripts were omitted from the analysis. Insertion counts were aggregated at the gene level. Background (input) replicates showed high reproducibility (Pearson correlation coefficient ≥ 0.895) and were therefore merged for subsequent analyses. To identify genes with significantly higher insertion counts in sorted samples compared to background, a one-sided Fisher’s exact test (alternative = ‘greater’) was performed on merged biological replicates, restricting the analysis to genes with at least 3 unique insertions in sorted samples. P values were adjusted for multiple testing using the false discovery rate (FDR) method.

To identify activators specific to CGI promoters, a stringent filtering strategy was applied. High-confidence hits were defined as genes that activated all 3 CGI reporters, using a strict statistical cutoff (FDR < 0.1%, log2 Odds Ratio ≥ 1). To ensure specificity, a comprehensive list of proteins capable of activating the non-CGI promoter was generated using a more lenient cutoff in the control screen (FDR < 5%, log2 Odds Ratio ≥ 1), and these were subtracted from the CGI hits.

### CUT&RUN analysis

Sequencing reads were trimmed to remove Illumina adapter sequences using Trim Galore (v0.6.0) with default parameters. The trimmed reads were aligned to the mm10 mouse genome assembly using Bowtie2 (v2.3.4.2, ^41^), with the maximum allowed insert size set to 1,000 bp. Reads with mapping quality scores ≤ 30 were removed using samtools (v1.9, ^42^). Peak identification was performed on individual and merged replicates using MACS2 (v2.1.2.1), with each sample compared against its respective input control. For paired-end libraries, the format was specified as BAMPE. For single-end libraries, model-free peak calling was performed with an extension size of 300 bp and a shift of 0. In all cases, duplicate reads were restricted to a single occurrence (--keep-dup 1), and normalized signal tracks (--SPMR) were generated. BedGraph files containing normalized coverage were converted to BigWig format using bedGraphToBigWig. Biological replicates exhibited high Pearson correlation and were therefore merged for subsequent analyses. For promoter assignment, only peaks within ±1,000 bp of gene transcription start sites (TSSs) were considered.

### Genome annotation

Gene annotation for the mm10 mouse genome was based on the GENCODE vM25 basic gene annotation. To define a single, representative transcription start site (TSS) for each gene, promoter coordinates were obtained from the Eukaryotic Promoter Database (EPDnew; v003 for coding and v001 for non-coding genes,^43^). For genes with multiple annotated promoters, the one with the highest CAGE (Cap Analysis of Gene Expression) signal, derived from FANTOM5 consortium data^44^, was selected as the dominant TSS. Promoter regions were defined as a 2 kb window centred on the TSS (±1 kb) and were annotated as CpG island (CGI) promoters if this region overlapped with experimentally defined CGI coordinates^45^, which were lifted over from the mm9 to the mm10 genome assembly.

### PRO-seq analysis

To remove PCR duplicates, a 10-nt unique molecular identifier (UMI) was incorporated at the 5′ end of reads during library preparation. UMIs were extracted and Illumina adapters trimmed with cutadapt (v1.18); reads shorter than 10 nt after trimming were discarded. For paired-end libraries only the first mate was used for alignment to the mouse mm10 genome with Bowtie (v1.2.2,^46^), allowing up to 2 mismatches and reporting a single best alignment per read (-m 1 --best --strata). To count unique nascent RNA molecules, reads that mapped to the same genomic coordinate were collapsed by UMI, permitting up to one mismatch within the UMI sequence. For PRO-seq coverage, only the first nucleotide of each read (representing the 3′ end of the nascent transcript) was recorded and strand information was adjusted to reflect transcription direction. Spike-in normalization was performed using reads aligned to the D. melanogaster (dm3) genome. For each sample, a size factor was calculated by dividing its individual UMI-corrected spike-in read count by the median spike-in read count calculated across all samples in the experiment. These manually calculated size factors were then used to normalize the endogenous gene counts within the DESeq2 analysis (v1.22.2,^47^). Genes were considered significantly differentially expressed if they met the criteria of an FDR < 0.05 and |fold change| > 1.5.

### Gene ontology term enrichment analysis

GO annotations for Biological Process, Molecular Function and Cellular Component were obtained from the org.Mm.eg.db R package (v3.17.0). Over-representation of GO terms among down-regulated genes was assessed by one-sided Fisher’s exact test (alternative = ‘greater’), using all expressed genes as the background set. P values were adjusted for multiple testing using the FDR method, and terms with FDR < 0.05 were considered significantly enriched. GO categories containing fewer than 5 genes in total or with fewer than 3 matching hits in the dataset were excluded. For visualization, the top 10 significantly enriched GO terms, ranked by FDR, were plotted.

### TurboID & IP-MS analysis

Differential protein abundance was determined using the limma R package (v 3.56.2). The analysis was performed on imputed and normalized apQuant Area values from three biological replicates, following the removal of common contaminants. Log2-transformed abundance values were fitted to a linear model, and empirical Bayes statistics were used to assess the significance of pairwise comparisons. P-values were adjusted using the FDR method, and high-confidence interactors were defined as proteins with a FDR < 0.05 and a log2FC > 1 relative to the control.

### Transcription factor motif analysis

Hcfc1-associated TFs were identified by filtering significant TurboID proteomics hits against a comprehensive TF annotation database^48^. A background set of all expressed TFs was compiled from PRO-seq data (baseMean > 5). Using a precomputed database of human TF motifs^49^, v2.1beta), the mean GC content was calculated for each position weight matrix (PWM). The preference for GC-rich motifs was assessed by comparing the GC content distribution of Hcfc1-associated TFs against all expressed TFs using a one-sided Wilcoxon rank-sum test (alternative = ‘greater’).

### AlphaFold2 Multimer screening

Protein interaction prediction screening was performed using a custom pipeline (HT-Colabfold) based on Colabfold, which utilizes AlphaFold2 Multimer^23,24,50^. This pipeline was used to perform two distinct sets of predictions: first, interactions between Hcfc1 domains and 127 proteins identified by TurboID; and second, interactions between the Hcfc1-AD and components of the pre-initiation complex (PIC). Predictions with an average iPTM score of > 0.5 or a PEAK score of > 0.75 (calculated as: PEAKavg = 1 - (max PAE in relevant quadrants / 31.75)) were considered putative hits and diagnostic plots (PAE plot, pLDDT plot and sequence coverage) as well as the generated structures were manually inspected^51^. All structural figures were prepared using UCSF ChimeraX^52^.

### Sequence searches and alignments

Sequences of target proteins were retrieved from the Uniprot database (https://www.uniprot.org) and aligned in Jalview using the integrated MUSCLE algorithm. Positional conservation and similarity scores were calculated using the ESPript webserver with default settings, and the resulting outputs were manually formatted to generate the final alignment figures.

## Data availability

All raw sequencing data have been deposited in the Gene Expression Omnibus (GEO) under accession **GSE307001** (accessible with secure token: mhcpoqoojtsvnmz). Mass spectrometry datasets have been deposited with the ProteomeXchange Consortium via the PRIDE repository under identifier **PXD067980** (accessible with secure token: vjWXpKMXGynp). Previously published datasets used for analysis were obtained from the following sources: non-CGI ORFtag data^14^, CpG island annotation^45^, transcription factors list^48^, and transcription factor binding motifs^49^. Source data are provided with the manuscript.

## Code availability

This study did not generate any original code. All analyses were performed using standard software as detailed in the Methods section.

## References

1. Haberle, V. & Stark, A. Eukaryotic core promoters and the functional basis of transcription initiation. Nat. Rev. Mol. Cell Biol. 19, 621–637 (2018).

2. Sainsbury, S., Bernecky, C. & Cramer, P. Structural basis of transcription initiation by RNA polymerase II. Nat. Rev. Mol. Cell Biol. 16, 129–143 (2015).

3. Carninci, P. et al. Genome-wide analysis of mammalian promoter architecture and evolution. Nat. Genet. 38, 626–635 (2006).

4. Deaton, A. M. & Bird, A. CpG islands and the regulation of transcription. Genes Dev. 25, 1010–1022 (2011).

5. Brent, R. & Ptashne, M. A eukaryotic transcriptional activator bearing the DNA specificity of a prokaryotic repressor. Cell 43, 729–736 (1985).

6. Haberle, V. et al. Transcriptional cofactors display specificity for distinct types of core promoters. Nature 570, 122–126 (2019).

7. Stampfel, G. et al. Transcriptional regulators form diverse groups with context-dependent regulatory functions. Nature 528, 147–151 (2015).

8. Freiman, R. N. & Herr, W. Viral mimicry: common mode of association with HCF by VP16 and the cellular protein LZIP. Genes Dev. 11, 3122–3127 (1997).

9. Wysocka, J., Reilly, P. T. & Herr, W. Loss of HCF-1-chromatin association precedes temperature-induced growth arrest of tsBN67 cells. Mol. Cell. Biol. 21, 3820–3829 (2001).

10. Dejosez, M. et al. Ronin/Hcf-1 binds to a hyperconserved enhancer element and regulates genes involved in the growth of embryonic stem cells. Genes Dev. 24, 1479–1484 (2010).

11. Thomas, L. R. et al. Interaction of MYC with host cell factor-1 is mediated by the evolutionarily conserved Myc box IV motif. Oncogene 35, 3613–3618 (2016).

12. Wysocka, J. & Herr, W. The herpes simplex virus VP16-induced complex: the makings of a regulatory switch. Trends Biochem. Sci. 28, 294–304 (2003).

13. Michaud, J. et al. HCFC1 is a common component of active human CpG-island promoters and coincides with ZNF143, THAP11, YY1, and GABP transcription factor occupancy. Genome Res. 23, 907–916 (2013).

14. Nemčko, F. et al. Proteome-scale tagging and functional screening in mammalian cells by ORFtag. Nat. Methods 1–6 (2024) doi:10.1038/s41592-024-02339-x.

15. Beerli, R. R., Segal, D. J., Dreier, B. & Barbas, C. F. Toward controlling gene expression at will: Specific regulation of the erbB-2/HER-2 promoter by using polydactyl zinc finger proteins constructed from modular building blocks. Proc. Natl. Acad. Sci. 95, 14628–14633 (1998).

16. Popay, T. M. et al. MYC regulates ribosome biogenesis and mitochondrial gene expression programs through its interaction with host cell factor-1. eLife 10, e60191 (2021).

17. Nishimura, K., Fukagawa, T., Takisawa, H., Kakimoto, T. & Kanemaki, M. An auxin-based degron system for the rapid depletion of proteins in nonplant cells. Nat. Methods 6, 917–922 (2009).

18. Arafeh, R., Shibue, T., Dempster, J. M., Hahn, W. C. & Vazquez, F. The present and future of the Cancer Dependency Map. Nat. Rev. Cancer 25, 59–73 (2025).

19. Goto, H. et al. A single-point mutation in HCF causes temperature-sensitive cell-cycle arrest and disrupts VP16 function. Genes Dev. 11, 726–737 (1997).

20. Van Nuland, R. et al. Quantitative Dissection and Stoichiometry Determination of the Human SET1/MLL Histone Methyltransferase Complexes. Mol. Cell. Biol. 33, 2067– 2077 (2013).

21. Sheikh, B. N., Guhathakurta, S. & Akhtar, A. The non-specific lethal ( NSL ) complex at the crossroads of transcriptional control and cellular homeostasis. EMBO Rep. 20, e47630 (2019).

22. Long, H. K., Blackledge, N. P. & Klose, R. J. ZF-CxxC domain-containing proteins, CpG islands and the chromatin connection. Biochem. Soc. Trans. 41, 727–740 (2013).

23. Evans, R. et al. Protein complex prediction with AlphaFold-Multimer. 2021.10.04.463034 Preprint at 10.1101/2021.10.04.463034 (2022).

24. Jumper, J. et al. Highly accurate protein structure prediction with AlphaFold. Nature 596, 583–589 (2021).

25. Luciano, R. L. & Wilson, A. C. An activation domain in the C-terminal subunit of HCF-1 is important for transactivation by VP16 and LZIP. Proc. Natl. Acad. Sci. 99, 13403–13408 (2002).

26. Patel, A. B., Greber, B. J. & Nogales, E. Recent insights into the structure of TFIID, its assembly, and its binding to core promoter. Curr. Opin. Struct. Biol. 61, 17–24 (2020).

27. Dikstein, R., Zhou, S. & Tjian, R. Human TAFII105 Is a Cell Type–Specific TFIID Subunit Related to hTAFII130. Cell 87, 137–146 (1996).

28. Falender, A. E. et al. Maintenance of spermatogenesis requires TAF4b, a gonad-specific subunit of TFIID. Genes Dev. 19, 794–803 (2005).

29. Gura, M. A. et al. Dynamic and regulated TAF gene expression during mouse embryonic germ cell development. PLOS Genet. 16, e1008515 (2020).

30. Chen, X. et al. Structures of +1 nucleosome–bound PIC-Mediator complex. Science 378, 62–68 (2022).

31. Warfield, L. et al. Transcription of Nearly All Yeast RNA Polymerase II-Transcribed Genes Is Dependent on Transcription Factor TFIID. Mol. Cell 68, 118–129.e5 (2017).

32. Sun, F. et al. The Pol II preinitiation complex (PIC) influences Mediator binding but not promoter–enhancer looping. Genes Dev. 35, 1175–1189 (2021).

33. Bergman, D. T. et al. Compatibility rules of human enhancer and promoter sequences. Nature 607, 176–184 (2022).

34. Liu, H. & Naismith, J. H. An efficient one-step site-directed deletion, insertion, single and multiple-site plasmid mutagenesis protocol. BMC Biotechnol. 8, 91 (2008).

35. Moussa, H. F. et al. Canonical PRC1 controls sequence-independent propagation of Polycomb-mediated gene silencing. Nat. Commun. 10, 1931 (2019).

36. Neumayr, C. et al. Differential cofactor dependencies define distinct types of human enhancers. Nature 606, 406–413 (2022).

37. Serebreni, L. et al. Functionally distinct promoter classes initiate transcription via different mechanisms reflected in focused versus dispersed initiation patterns. EMBO J. 42, e113519 (2023).

38. Yusa, K., Zhou, L., Li, M. A., Bradley, A. & Craig, N. L. A hyperactive piggyBac transposase for mammalian applications. Proc. Natl. Acad. Sci. 108, 1531–1536 (2011).

39. Skene, P. J. & Henikoff, S. An efficient targeted nuclease strategy for high-resolution mapping of DNA binding sites. eLife 6, e21856 (2017).

40. Lawrence, M. et al. Software for computing and annotating genomic ranges. PLoS Comput. Biol. 9, e1003118 (2013).

41. Langmead, B. & Salzberg, S. L. Fast gapped-read alignment with Bowtie 2. Nat. Methods 9, 357–359 (2012).

42. Danecek, P. et al. Twelve years of SAMtools and BCFtools. GigaScience 10, giab008 (2021).

43. Dreos, R., Ambrosini, G., Groux, R., Cavin Périer, R. & Bucher, P. The eukaryotic promoter database in its 30th year: focus on non-vertebrate organisms. Nucleic Acids Res. 45, D51–D55 (2017).

44. Forrest, A. R. R. et al. A promoter-level mammalian expression atlas. Nature 507, 462–470 (2014).

45. Illingworth, R. S. et al. Orphan CpG islands identify numerous conserved promoters in the mammalian genome. PLoS Genet. 6, e1001134 (2010).

46. Langmead, B., Trapnell, C., Pop, M. & Salzberg, S. L. Ultrafast and memory-efficient alignment of short DNA sequences to the human genome. Genome Biol. 10, R25 (2009).

47. Love, M. I., Huber, W. & Anders, S. Moderated estimation of fold change and dispersion for RNA-seq data with DESeq2. Genome Biol. 15, 550 (2014).

48. Lambert, S. A. et al. The Human Transcription Factors. Cell 172, 650–665 (2018).

49. Vierstra, J. et al. Global reference mapping of human transcription factor footprints. Nature 583, 729–736 (2020).

50. Mirdita, M. et al. ColabFold: making protein folding accessible to all. Nat. Methods 19, 679–682 (2022).

51. Hohmann, U. et al. A molecular switch orchestrates the nuclear export of human messenger RNA. 2024.03.24.586400 Preprint at 10.1101/2024.03.24.586400 (2024).

52. Goddard, T. D. et al. UCSF ChimeraX: Meeting modern challenges in visualization and analysis. Protein Sci. Publ. Protein Soc. 27, 14–25 (2018).

